# Motor Cortical Output Integrates Distorted Proprioceptive Feedback

**DOI:** 10.1101/2025.11.11.687832

**Authors:** Yi Li, X. Hermione Xu, Shiyang Pan, Xu An, Patrick J. Mulcahey, Zixuan Qiu, Shengli Zhao, Nuo Li, John Pearson, Z. Josh Huang

## Abstract

Proprioceptive feedback from muscles is essential for continuous monitoring and precise control of limb movement, yet how such peripheral feedback is integrated into ongoing descending motor cortical dynamics remains largely unclear. Here we used precisely timed optogenetic stimulation of forelimb muscles to manipulate proprioceptive signals during a mouse reach-to-consume task. Mice were able to successfully reach and consume despite muscle stimulation-evoked forelimb deviations. Across the cortex, the caudal forelimb area (CFA) was preferentially activated by muscle-specific proprioceptive inputs and contributed to stabilizing perturbed forelimb movement. CFA extratelencephalic (ET) neurons encoded movement kinematics as well as proprioceptive feedback. At neuron population level, proprioceptive perturbations rapidly deflected CFA ET neural trajectories away from the task-relevant manifold yet exerted only limited effects on evolving task dynamics. These results reveal that descending cortical circuit implements activity subspace separation to encode and integrate distorted proprioceptive feedback while preserving task-relevant output.

## Introduction

Motor systems in animals rely on mechanosensory feedback from the somatosensory system for optimal control and precision ^1,2^. Among mechanosensory modalities, proprioceptive feedback arises from muscle and joint receptors, allowing the nervous system to continuously estimate movement position, velocity and force ^3–5^. The brain monitors sensory feedback and compares these signals with internal predictions to achieve skilled control of motor behavior ^6–8^. Long-term loss of proprioceptive input from lesions or disease compromises movement precision, coordination and adaptability ^9–12^.

At the neural circuit level, proprioceptors in dorsal root ganglia (DRG) detect muscle stretch and contraction and project to the spinal cord and brainstem circuits ^13^. From there, signals ascend to the forebrain primarily through the dorsal column-medial lemniscal pathway via thalamic relay nuclei to sensorimotor cortex. In the cortex, ascending proprioceptive signals converge with descending motor commands ^14–16^. Motor cortical neurons respond to both voluntary movement and passive limb displacement ^14–21^. Such feedback integration supports online corrections ^1,16,21,22^ and calibrates internal models that predict sensory consequences of action ^23,24^. Artificial sensory feedback can improve operant control and learning ^25,26^ and persistent forelimb perturbations have demonstrated feedback-driven adaptation ^27–29^. Proprioceptive feedback from muscles therefore enables the cortex to continuously monitor and precisely control limb movement. As proprioceptive signals are transmitted by multiple receptor systems, it has been challenging to precisely manipulate such feedback from defined muscles during complex movement while simultaneously monitoring cortical activity. Thus, how motor cortex processes muscle-specific proprioceptive inputs during movement remains poorly understood.

Neural population dynamics in motor cortex encode motor plans and kinematics ^30–33^. Descending cortical output, carried by extratelencephalic (ET) neurons that broadcast signals to thalamic, striatal, brainstem, and spinal circuits ^34–36^, is essential for controlling ongoing self-initiated forelimb movement ^37^. ET outputs and thalamic inputs form a closed-loop circuit ^38^, in which thalamic integration of ongoing sensorimotor state shapes the evolution of cortical dynamics ^37,39^. Limb perturbations alter somatosensory inflow. Expectations about such disturbances shape preparatory dynamics ^40^, and can index learned motor memories ^41^. Beyond compensation based on expectation or memory, a stable controller must preserve descending dynamics for movement generation while integrating altered sensory feedback. How, then, does the same population of cortical output neurons process distorted feedback without substantially corrupting ongoing behavior? One proposed mechanism is that cortical populations implement distinct computations by partitioning activity into separable subspaces ^32,33,42^, potentially segregating feedback-related signals from task-defining output. However, it remains unclear how cortical output circuits actually integrate natural and distorted proprioceptive feedback into ongoing population activity during motor control.

To address this gap, we measured cortical output population dynamics while selectively and rapidly activating proprioceptive signals during a mouse reach-to-consume task. Using optogenetic muscle stimulation, we achieved precisely timed manipulation of forelimb movement and proprioceptive signals. While muscle stimulation reliably deflected forelimb kinematic trajectories, mice were still able to successfully reach and consume. Across cortical areas, the CFA preferentially processed proprioceptive signals from forelimb muscles and contributed to stabilizing forelimb movement. Combining proprioceptive perturbation with simultaneous two-photon imaging of CFA ET neuronal activity, we found that motor cortical output populations route distorted proprioceptive signals into subspaces largely separate from the task-relevant manifold, preserving ongoing descending dynamics. These results identify subspace separation as a neural mechanism through which descending motor cortical dynamics rapidly integrate proprioceptive feedback while maintaining task-relevant output.

## Results

### Selective Manipulation of Proprioceptive Feedback by Optogenetic Muscle Stimulation

Selectively manipulate proprioceptive inputs from specific muscle effectors is necessary to causally identify feedback signals and address how it shapes cortical dynamics. Proprioceptive inputs from muscle spindles and Golgi tendon organs detect changes in muscle length and force and relay this information to the spinal cord and brain via group Ia and Ib sensory neurons in the dorsal root ganglia (**Fig. 1A**). To manipulate these signals from the forelimb with precise temporal control and identify feedback activity to the cortex, we optogenetically activated skeletal muscles using a biocompatible implant. To express ChR2 in muscle cells, we packaged the channelrhodopsin (ChR2) gene into a muscle-enriched AAV serotype (AAVMYO3 ^43^) and locally injected the viral vector into the left biceps and triceps (**Fig. 1B**). Immunohistochemistry confirmed robust ChR2 expression in the sarcolemma of muscle fibers (**Fig. 1B, S1A**). To precisely deliver light pulses to the muscles across behavioral contexts, we engineered a biocompatible mini-LED and implanted it subcutaneously in the peripheral forelimb with the connector fixed on the skull (**Fig. S1B**).

**Fig 1.**
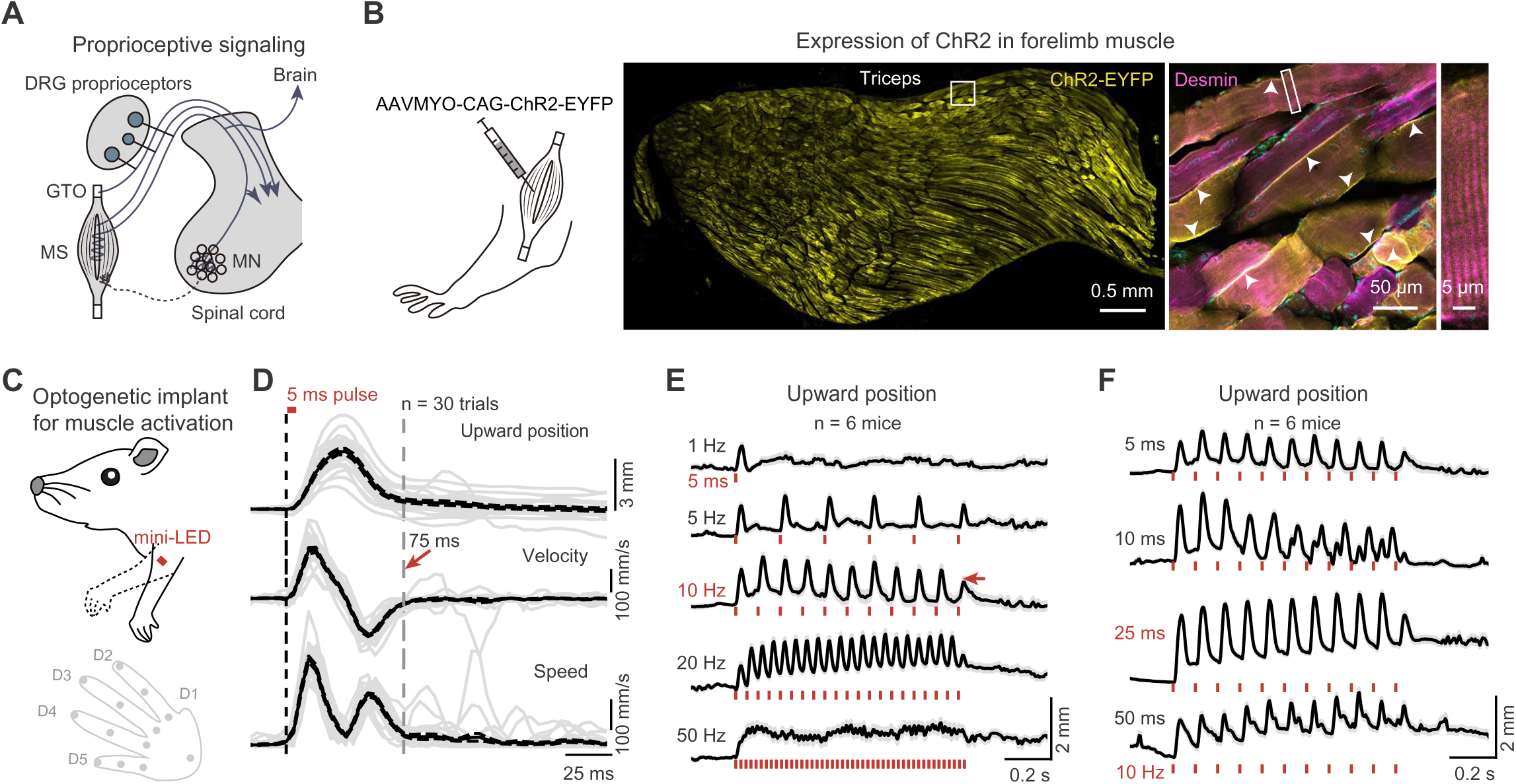
Manipulation of forelimb proprioceptive signals by optogenetic stimulation of muscles. **(A)** Peripheral components transmitting movement feedback to the spinal cord and brain. MS, muscle spindles; GTO, Golgi tendon organ; DRG, dorsal root ganglion; MN, motor neurons. **(B)** Myotropic adeno-associated virus (AAVMYO) expression of ChR2 in mouse forelimb muscles. Desmin staining outlines the sarcomere boundaries. Arrows indicate sarcolemma membrane expression. **(C)** Top: Implantation of a mini-LED on the forelimb proximal muscles for optogenetic stimulation. Bottom: Tracking of digits (D) to quantify forelimb movement. **(D)** Hand movement kinematics evoked by a single 5 ms light pulse from a session (n = 30 trials). **(E)** Upward hand movement evoked by 1 s trains of 5 ms light pulses at 1, 5, 10, 20, and 50 Hz (n = 6 mice, mean ± SEM). The reduced amplitude (arrow) results from the final pulse being missed in some mice. **(F)** Upward hand movement evoked by trains of 5, 10, 25, and 50 ms light pulses at 10 Hz (n = 6 mice, mean ± SEM). Note the larger amplitude and minimal decay across repeated 25 ms stimulation pulses.

A single brief optogenetic light pulse triggers a forelimb movement cycle consisting of a contraction-evoked displacement followed by a return (**Supplementary Video 1**). ChR2-expressing and LED-implanted mice were head-restrained with their forelimbs hanging freely and relaxed (**Fig. 1C**). To quantify limb movements, we trained deep neural networks to extract key points on the left hand from high-speed front- and side-view videos. Left-hand kinematics were computed and aligned to the onset of light pulses. Brief 5-ms optogenetic pulses (40 mW) reliably displaced the forelimb with a peak speed of ∼30 cm/s within 30 ms of light onset (**Fig. 1D**). After reaching maximum displacement, the forelimb typically returned to its starting position with a slower peak speed (∼20 cm/s) (**Fig. 1D, S1C**). The return is unlikely due to a simple vertical gravity effect as the limb displacement along the forward direction also returns (**Fig. S1D**). The total duration of this displacement-return cycle was approximately 75 ms, beyond which we did not observe reliable limb movement (**Fig. 1D, S1E**). This rapid sequence of muscle contraction and limb movement likely alters Group Ia and Ib proprioceptive inputs, which are sensitive to dynamic changes in muscle length and tension, respectively.

A train of light pulses reliably activated muscles and evoked pulse-locked oscillatory movements of the forelimb (**Supplementary Video 1**). Across stimulation frequencies ranging from 1 to 50 Hz, 10 Hz pulses produced robust limb displacements while preserving the full displacement-return cycle (**Fig. 1E, S1F**). We further tested a range of pulse durations from 5 ms to 50 ms and found that 25 ms pulses elicited the largest amplitude of forelimb displacement (**Fig. 1F**). Progressively longer duration stimulations displace the limb without a full return coupled to each pulse (**Fig. 1F**). Based on these data, we used 25 ms light pulses at 10 Hz to effectively actuate muscle contraction as a method to generate precisely timed proprioceptive inputs to the brain.

Consistent with direct muscle activation, optogenetic stimulation continued to evoke displace-return movement even in the spinal-lesioned limbs of anesthetized mice (**Fig. S1G**), confirming that the passively elicited movement was primarily muscle contraction-relaxation cycle derived and not dependent on spinal reflexes.

In summary, we have successfully developed and validated a reliable **opto**genetic strategy to selectively manipulate **f**orelimb **p**roprioceptive feedback by activating muscle contractions using a biocompatible mini-LED implant (**optoFP**). This strategy allows us to precisely manipulate proprioceptive feedback signals and reveal how it shapes cortical dynamics.

### Reach-to-consume Deviations Under Transient Proprioceptive Perturbations

Optogenetic stimulation of forelimb muscles rarely blocked reach-to-consume progression in head-fixed mice (**Supplementary Video 2**). We trained mice to retrieve water droplets from a fixed waterspout using their left forelimbs ^37,44^. In perturbation trials, we applied 25 ms light pulses at 10 Hz for 2 s, triggered around reach onset, to perturb forelimb reach-to-consume action (**Fig. 2A**). We employed a 40 baseline / 60 perturbation / 40 recovery block design in each session. Across 27 sessions (2,160 control trials), mice initiated forelimb movement 0.37 ± 0.59 s (mean ± SD; n = 2120/2160 trials) after water delivery, completed reach, grasp and withdraw in 0.49 ± 0.69 s, followed by 2.23 ± 1.35 s of hand-assisted licking, consistent with our prior results ^37^. In perturbed trials, the first stimulation pulse occurred 0.10 ± 0.32 s after reach onset (n = 1,582/1,620; **Fig S2A**). Over half of successful grasp trials (872/1,554) received fewer than two pulses during rapid reaches (**Fig. S2B**). Most stimulation pulses were after grasp during hand-assisted consumption (**Fig. S2B**). Despite evoked muscle contractions, optogenetic forelimb stimulation did not significantly reduce overall grasp success rates (Ctrl 1971/2160 *vs* Pert 1555/1620), though it slightly delayed grasp latency (**Fig. 2B**, Ctrl 0.25 ± 0.02 s *vs* Pert 0.29 ± 0.02 s). These findings indicate that mice were able to successfully control movement even in the presence of transient peripheral perturbations.

**Fig 2.**
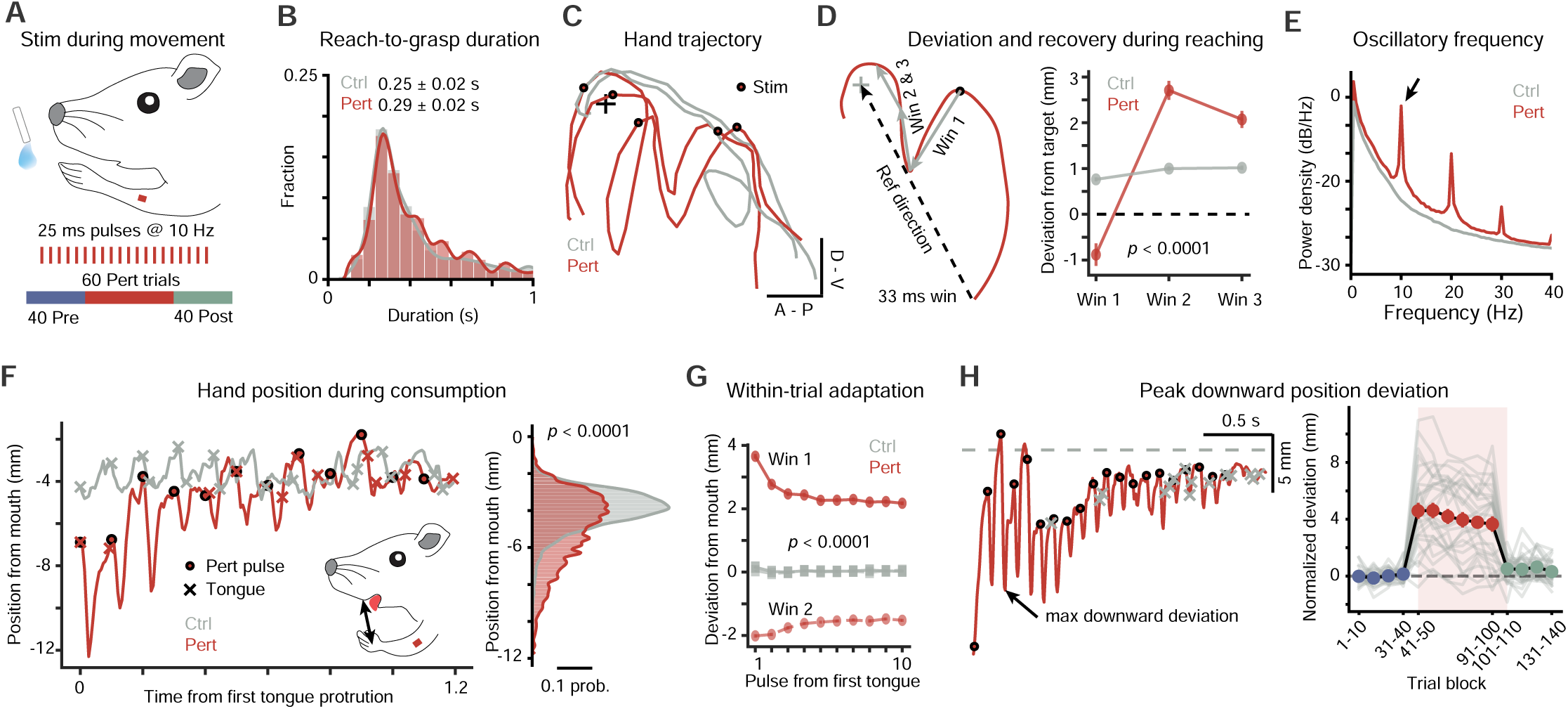
Optogenetic muscle stimulation deflects reach-to-consume movement. **(A)** Optogenetic forelimb muscle stimulation during reach-to-consume actions. The waterspout was fixed. Each session included 40 pre-perturbation baseline (Pre), 60 perturbation (Pert), and 40 post-perturbation recovery (Post) trials. Light pulses (25 ms, 10 Hz) were triggered for 2 s around reach onset. **(B)** Distribution of reach-to-grasp durations from control (Ctrl) and perturbation (Pert) trials (Wilcoxon rank-sum test, p = 0.0002; n = 1971 Ctrl and 1555 Pert trials from 27 sessions, 4 mice). **(C)** Example hand trajectories from two Ctrl and two Pert trials. **(D)** Left: Schematic of movement deviation analysis. Three consecutive 33 ms trajectory fragments were projected onto the reference direction. In Ctrl trials, “virtual” light pulses were used for alignment. Right: Single-pulse forelimb stimulations induced deviations away from the target in the first window, followed by recovery (n = 2160 Ctrl trials and 163 Pert trials with single-pulse perturbations). **(E)** Power spectral density of forelimb trajectories (n = 2160 Ctrl and 1620 Pert trials). Arrow indicates the perturbation-coupled 10 Hz oscillatory frequency. **(F)** Left: Hand position relative to the mouth during water consumption in a Ctrl and a Pert trial. Right: Distribution of hand position upon tongue protrusions with and without forelimb stimulations (Wilcoxon rank-sum test, p < 0.0001; n = 28,870 Ctrl and 7830 Pert tongue protrusions). **(G)** Within-trial decay in perturbation-evoked peak downward hand deviations from the mouth across stimulation pulses during consumption (two-way ANOVA for each 33 ms window: Group, p < 0.0001; Pulse sequence, p < 0.0001; Group × Pulse interaction, p < 0.0001; n = 1620 Ctrl and 1620 Pert trials; mean ± SEM). **(H)** Left: Identification of peak downward deviation of hand position. Right: Peak downward deviations across trials from each session (n = 27 sessions; mean ± SEM). Deviations were averaged per 10 trials and baseline-subtracted for each session. Note the slight decrease trend across Pert trials.

Transient forelimb stimulations briefly deviated mouse reaching kinematics (**Supplementary Video 2**). Unlike the stable trajectories in control trials, perturbed reaches showed brief, high-speed deviations (∼35 cm/s) away from the target within 33 ms of stimulation pulse onset (**Fig. 2C-D**). In the subsequent 33 ms before the next pulse, the downward-deviated hand typically reoriented toward the target instead of bouncing back directly to its pre-deviation position given the inertia of limb motion (**Fig. 2C-D**). No such deviation-recovery cycles were observed in control trials in which “virtual” stimulation pulses were assigned to randomly selected windows of equivalent duration. The pattern of deviation followed by partial return produced oscillatory trajectories at 10 Hz, 20 Hz and harmonic frequencies (**Fig. 2E**), yet overall hand movement continues to progress toward the target.

Downward forelimb deviations decreased during hand-assisted consumption (**Supplementary Video 2**). Optogenetically induced deviations varied with stimulation timing relative to the reach-to-consume sequence, with amplitude increasing as the hand approached the spout and decreasing post-grasp (**Fig. S2C-D**). In control trials, post-grasp hand withdrawal stabilized near the mouth with minor oscillations (**Fig. 2F**). Under optoFP, the hand initially deflected downward post-grasp but was progressively guided back toward the mouth, closely matching control trial positions during licking (**Fig. 2F**). The amplitude of the evoked downward deviations and upward recovery per light pulse were significantly larger in perturbation trials than in virtual pulses as control (**Fig. 2G**). Across the stimulation pulses during consumption, the downward position deviation amplitude exponentially decreased (**Fig. 2G**). This attenuation is better explained by increasing limb resistance as posture stabilizes than by pulse habituation, because passive stimulation outside behavior produced stable deviations and repeated pulses during reach did not reduce deviation amplitude. Together, these observations indicate rapid within-trial stabilization, likely mediated by progressively enhanced limb stiffness during hand-assisted consumption.

Within the perturbation block, downward deviation amplitude showed only slight linear decrease across trials, indicating modest adaptation (-0.02 mm/trial, linear regression R^2^ = 0.9461; p = 0.0011; **Fig. 2H**). When the perturbation was unexpectedly removed in subsequent trials, the downward deviations rapidly decreased to baseline level (**Fig. 2H**). To assess adaptation across all kinematic dimensions, we defined a Behavior Discriminant Axis (BDA) by extracting behavioral latent via principal component analysis (PCA) and applying linear discriminant analysis to optimally separate perturbed from unperturbed trials (**Fig. S2E**). Projection onto the BDA confirmed the reversible increase in perturbation effects with only slight within-block adaptation (-0.0101 au/ trial, p = 0.0005; R^2^ = 0.9628; **Fig. S2E**). Removal of perturbation led to a rapid return to baseline without overshoot, indicating that muscle stimulations did not induce classical aftereffects under force-field perturbations^27,29^. Meanwhile, oscillatory energy slightly decayed across perturbation trials (p = 0.0405; R^2^ = 0.6904; **Fig. S2F**), whereas evoked hand speed remained unchanged (p = 0.6977; **Fig. S2G**), suggesting that across-trial adaptation may be specific to task-relevant kinematic features.

### Proprioceptive Signals Preferentially Activate the Caudal Forelimb Area

To map the cortical regions activated by forelimb proprioceptive input, we used wide-field calcium imaging to measure cortical responses to optogenetic forelimb muscle stimulation in head-fixed mice at rest (**Fig. 3A, S3A**). Specifically, we crossed *Slc17a7-ires-Cre* driver mice with GCaMP reporter lines (*GCaMP6f*) to monitor projection neuron activity (**Fig. S3B**). These mice also received *AAVMYO-ChR2-EYFP* injection into the forelimb muscle and mini-LED implantation for muscle activation. Optogenetic stimulation (10 Hz, 1 s) of the forelimb muscle robustly increased calcium fluorescence in the contralateral hemisphere, imaged through the intact transparent skull. These signals originated in a localized somatic region of the posterior and lateral caudal forelimb area (CFA) and spreading to broader regions before returning to its initial site (**Fig. 3A, S3C, Supplementary Video 3**). The right hemisphere CFA showed significantly higher activity than the left, indicating selective responsiveness to left forelimb stimulation (**Fig. 3B**). Calcium activity also rose in the rostral area and other regions (**Fig. 3C**). Within CFA, passive muscle activation evoked highly reliable calcium responses across trials (**Fig. 3D**). CFA is known to be enriched with corticospinal output neurons ^36^ and direct optogenetic activation of CFA ET neurons evoked robust forelimb movements ^45^ (**Fig. S3D-F**). These findings demonstrate that forelimb muscle stimulation preferentially activates proprioceptive signal processing in a forelimb movement-related cortical region.

**Fig 3.**
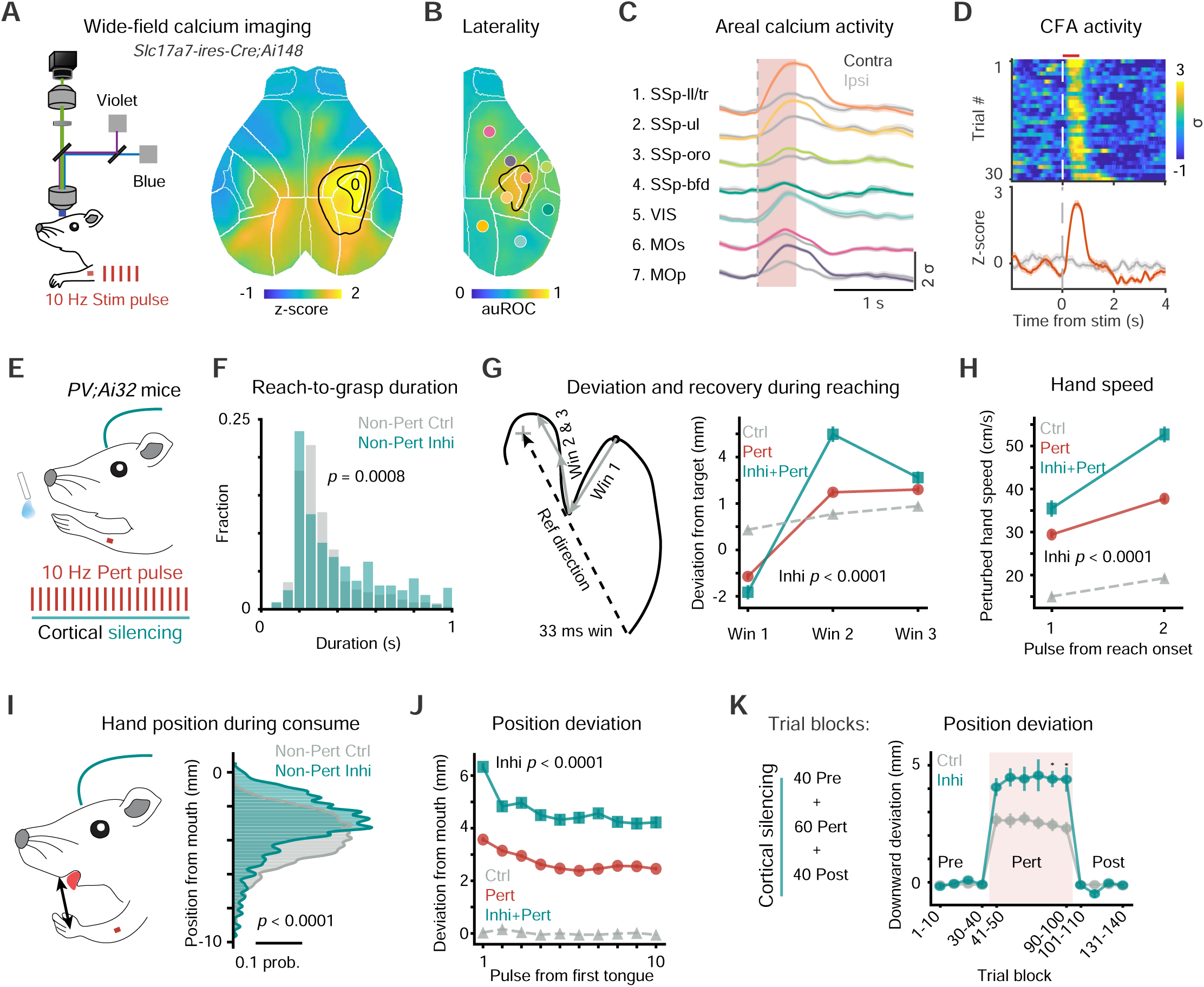
CFA inactivation exacerbates evoked forelimb deviations during the reach-to-consume task. **(A)** Left: Schematic for wide-field calcium imaging of cortical dynamics during passive forelimb muscle stimulation. Right: Average imaging frame showing strong calcium activity in the caudal forelimb area (CFA) evoked by passive left forelimb muscle stimulation (10 Hz, 0.5 s; n = 6 sessions) in *Slc17a7-ires-Cre;Ai148* mice. **(B)** Hemisphere laterality scores (auROC) of passively evoked calcium activity (n = 6 sessions). Larger auROC values indicate higher calcium activity in the right vs. left hemisphere. **(C)** Average calcium traces across cortical regions aligned to passive stimulation onset (n = 30 trials; mean ± SEM). **(D)** Heatmap and line plot of CFA calcium activity to passive muscle stimulation across trials (n = 30 trials). Trials without forelimb stimulations were used as controls (n = 30 trials). **(E)** Inactivating CFA under normal and perturbed reach-to-consume actions in *PV;Ai32* mouse. **(F)** Slightly changed distribution of grasp latency from reach onset from unperturbed control (Ctrl) and CFA inhibition (Inhi) trials (Wilcoxon rank-sum test, p = 0.0008; n = 990 Ctrl and 315 Inhi trials from 14 Ctrl and 4 Inhi sessions, 4 mice). **(G)** Left: Schematic of movement deviation analysis across three consecutive time windows for each stimulation pulse during reaching. Right: Inactivating CFA exacerbates initial deviations away from the target and the following recovery (two-way ANOVA for each window: Group [Pert vs Inhi+Pert], p < 0.0001; Pulse sequence, p < 0.0001; Group × Pulse interaction, p < 0.0001; n = 729 Ctrl, 491 Pert and 142 Inhi+Pert trials with single-pulse stimulations; mean ± SEM). **(H)** Increase in peak deviation speed of the two stimulation pulses during reaching two-way ANOVA for each window: Group [Ctrl vs Pert], p < 0.0001; Pulse sequence, p < 0.0001; Group × Pulse interaction, p < 0.0001; n = 1120 Ctrl, 840 Pert and 240 Inhi-Pert trials with single-pulse stimulations; mean ± SEM. **(I)** Left: Hand position upon tongue protrusions during consumption. Right: Cortical silencing altered the distribution of hand positions upon tongue protrusions of unperturbed consumption (Wilcoxon rank-sum test, p < 0.0001; n = 11,078 Ctrl and 2043 Inhi tongue protrusions). **(J)** Downward position deviations from mouth across successive stimulation pulses during hand licking (two-way ANOVA: Group [Pert vs Inhi+Pert], p < 0.0001; Pulse sequence, p < 0.0001; Group × Pulse interaction, p < 0.0001; n = 1120 Ctrl, 840 Pert and 240 Inhi+Pert trials with single-pulse stimulations; mean ± SEM). **(K)** Left: Cortical silencing across Pre, Pert and Post blocks in each session. Right: Critical silencing on downward deviations across trials from Ctrl and Inhi sessions (two-way ANOVA: Group [Ctrl *vs* Inhi], p < 0.0001; Trial block, p < 0.0001; Group × Trial interaction, p < 0.0001; Bonferroni post hoc test; n = 14 Ctrl and 4 Inhi sessions; mean ± SEM). Deviations were averaged per 10 trials and baseline-subtracted for each session.

### Inactivating CFA Exacerbates Evoked Forelimb Deviations

We hypothesized that CFA activity might be required to counteract muscle stimulation-evoked forelimb deviations. To test this, we optogenetically silenced CFA by activating cortical parvalbumin (PV) interneurons in *PV;Ai32* mice during forelimb muscle stimulation (**Fig. 3E**).

In addition to the expected cortical expression, PV is also expressed in skeletal muscle ^46^. Accordingly, *PV;Ai32* mice showed strong ChR2-EYFP labeling in forelimb muscle fibers, particularly at the sarcolemma (**Fig. S4A**). Although PV is present in subsets of dorsal root ganglion neurons and can label peripheral proprioceptive nerve terminals ^47^, we did not detect robust fluorescence in triceps or biceps nerves after staining, likely reflecting low ChR2-EYFP level at peripheral nerve terminals in these mice (**Fig. S4A**). Brief optogenetic stimulation through the forelimb implant (5 ms) evoked pulse-locked movements with short latency across a broad frequency range (**Fig. S4B-C**). These movements persisted even when stimulating the spinal-lesioned or excised forelimb in anesthetized *PV;Ai32* mice, consistent with direct activation of muscle rather than modulation of peripheral nerves (**Fig. S4D**). In awake, resting *PV;Ai32* mice, inactivating CFA did not significantly alter the kinematics of passively evoked forelimb movements (**Fig. S4E-F**), indicating minimal central modulation of muscle-stimulation-evoked forelimb movement under passive conditions.

We next asked whether and how CFA inactivation might modulate optoFP-evoked forelimb deviations during voluntary reach-to-consume behavior. In unperturbed trials, inactivating CFA alone rarely affected grasp success and produced only a small increase in grasp latency from reach onset (**Fig. 3F**). In perturbed trials, however, inactivating CFA increased both the deviation magnitude away from the target and the subsequent recovery amplitude during the reach (**Fig. 3G**). This exacerbation was larger on the second stimulation pulse, with higher peak deviation speed than after the first pulse (**Fig. 3H**).

During hand-assisted consumption phase, CFA inactivation exacerbated optoFP-evoked deviations. In unperturbed trials, it shifted hand position closer to the mouth during tongue protrusions, consistent with reduced positional precision (**Fig. 3I**). Under perturbation, CFA inactivation produced larger downward deviations and greater upward recovery across stimulation pulses (**Fig. 3J**), and increased optoFP-evoked peak speeds (**Fig. S4G**). These results indicate that ongoing CFA dynamics are required to counteract perturbation-evoked deviations and stabilize hand position during reach-to-consume behavior.

We then examined effects of CFA inactivation on the decrease of perturbation-evoked deviations. During consumption, within-trial decay of deviation amplitude across successive pulses was only slightly altered (**Fig. 3J, S4G**). Across perturbation trials, CFA silencing abolished the modest decay of downward position deviations seen under control conditions (**Fig. 3K**), yet the oscillatory energy of the evoked response still declined over pulses (**Fig. S4H**). Notably, CFA silencing alone also significantly altered oscillatory energy in unperturbed trials (**Fig. S4H**). Two-factor analyses revealed a significant interaction between forelimb stimulation and CFA inactivation for oscillatory energy (**Fig. S4H**), suggesting that the exacerbation reflects combined effects of peripheral perturbation and cortical inhibition. Overall, the impact of CFA inactivation on deviation decay was modest and appeared to be specific to task parameters.

Collectively, the exacerbated deviations hand position induced by optoFP during CFA silencing support a key role for CFA dynamics in stabilizing perturbed reach-to-consume actions.

### CFA ET Neurons Encode Natural and Distorted Proprioceptive Feedback

Given our findings that optogenetic muscle stimulations evoke activity in CFA, we hypothesized that such distorted proprioceptive feedback would modulate the dynamics of ET projection neurons there, potentially supporting motor stabilization under perturbation. To test this, we performed two-photon calcium imaging of CFA ET neurons in *Fezf2-CreER;GCaMP8m* mice (**Fig. 4A**). The same animals received viral ChR2 expression in forelimb muscles for optogenetic stimulation (**Fig. 4A**). GCaMP-labeled somata were enriched in cortical deep layers 5/6 ET projection neurons ^34^, projecting to the striatum, thalamus, brainstem, and spinal cord (**Fig. S5A**). After applying quality-control criteria, we analyzed calcium signals from 558 layer-5 ET neurons in CFA (**Fig. 4B-C, S5B**). Among them, 71.2% of CFA ET neurons (397/558) were significantly modulated during the reach-to-consume action (**Fig. 4D-E**).

**Fig 4.**
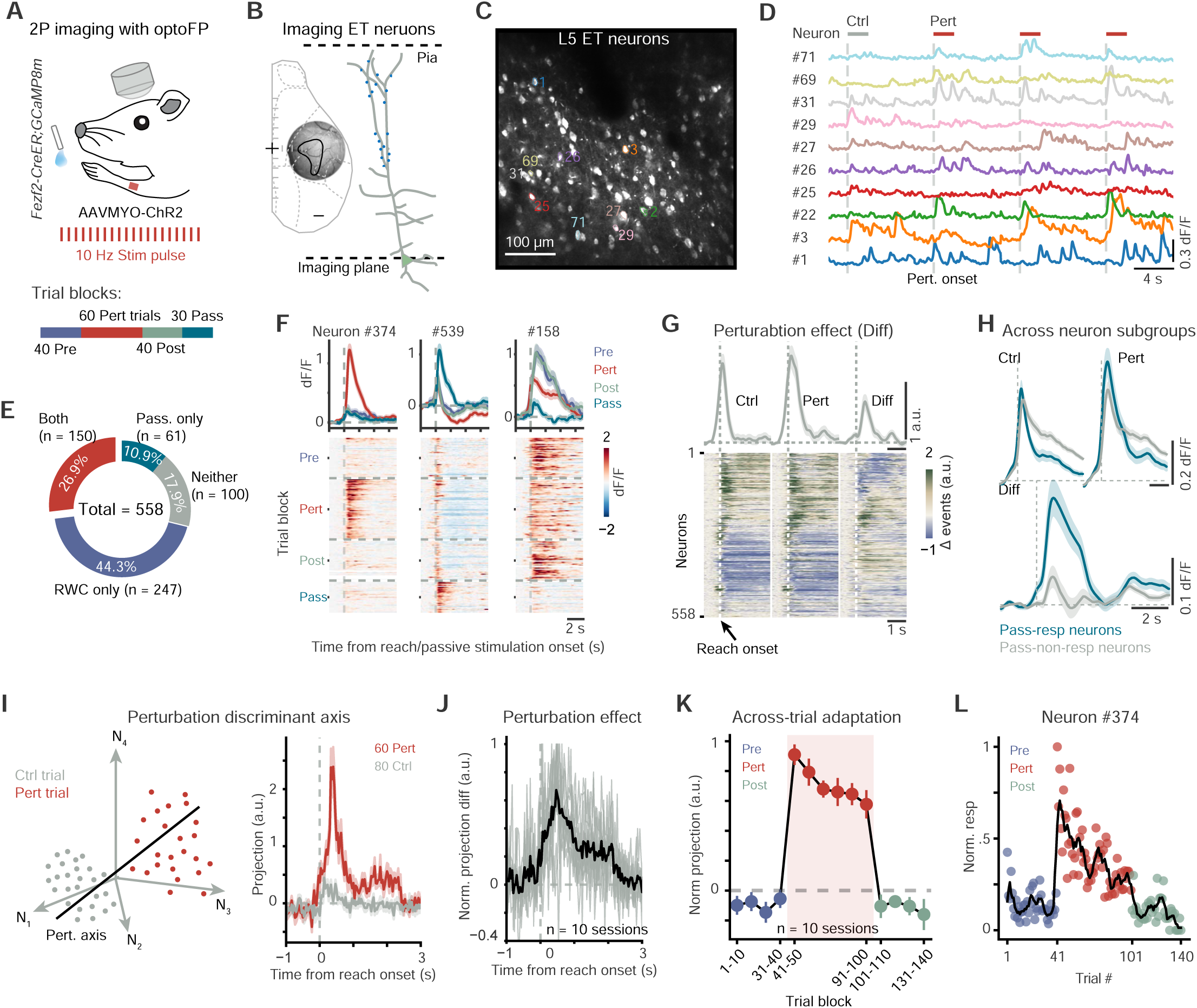
Normal and distorted proprioceptive signals in CFA extratelencephalic neurons. **(A)** Two-photon (2P) calcium imaging of CFA extratelencephalic (ET) neurons under forelimb stimulation in *Fezf2-CreER;GCaMP8m* mice. Each session consisted of 40 pre-perturbation (Pre), 60 perturbation (Pert), 40 post-perturbation (Post), and 31 passive muscle stimulation (Pass) trials. The stimulation consisted of 25 ms light pulses at 10 Hz for 2 s. **(B)** Left: Imaging window in CFA. Right: Schematic showing the imaging plain at layer 5 ET neuron. **(C)** A session-averaged imaging frame for ET neurons. **(D)** Activity traces from control and perturbed reach-to-consume trials. Vertical dashed lines indicate actual stimulation onset in perturbation trials or virtual stimulation onset in control trials. **(E)** Fraction of neurons modulated by reach-to-consume, passive muscle activation, or both. **(F)** Line plot summary and heatmap showing individual neuron activity across trials (n = 40 Pre, 60 Pert, 40 Post, and 30 passive stimulation [Pass] trials; mean ± SEM). Activity was aligned to voluntary reach onset or to passive forelimb stimulation onset. **(G)** Perturbation effect after deconvolution across all neurons (n = 558 neurons). Activity aligned at reach onset. Pre and Post trials were combined as control (Ctrl). **(H)** Larger perturbation effects in passive responsive neurons than that of passive-non-responsive neurons (n = 211 and 347 neurons; mean ± SEM). **(I)** Left: Schematic showing identification of the perturbation discriminant axis (PDA) in multidimensional neural activity space. Each dot represents the activity during the perturbation period of an individual trial; virtual stimulation windows were used for control trials. Right: Projected activity along the PDA from an example session (n = 60 Pert and 80 Ctrl trials; mean ± SEM). **(J)** Summary of the perturbation effect along the PDA across sessions (n = 10 sessions). Perturbation effect is the difference between PDA-projected activity in Pert and control trials and peak-normalized per session. **(K)** Summary of PDA-projected activity across trials (n = 10 sessions; mean ± SEM); activity was averaged per 10 trials and peak-normalized per session. **(L)** Example neuron showing decreased activity across Pert trials. Each dot represents neural activity during the perturbation period of one trial. The black line is the smoothed trace (sliding window). Activity was quantified as spike-event counts and normalized to the peak.

To quantify perturbation effects at the single-neuron level, we compared ET neuron activity between optogenetically perturbed versus unperturbed (virtual) reach trials (**Fig. 4F**). Fluorescent calcium signals were deconvolved to estimate relative change of spike events (**Fig. S5C**). More than half of CFA ET neurons exhibited significant, heterogeneous perturbation effects (56.6%, 316/558; **Fig. 4G**). On average, perturbation evoked a net increase in ET neuron activity relative to unperturbed reach-to-consume (**Fig. 4G**). The amplitude and peak latency of perturbation effects varied between neurons, typically peaking ∼0.5 s after reach onset before the stimulation ended (**Fig. 4G, S5D**), coinciding with the hand-assisted consumption. The amplitude of perturbation effect correlated with reach response amplitude (R^2^ = 0.395, **Fig. S5E**). These single-cell perturbation effects suggest CFA ET output populations encode distorted proprioceptive feedback.

We then asked whether CFA ET neurons encode both natural and distorted proprioceptive feedback during reach-to-consume behavior. Responses to passive muscle stimulation in resting mice indicate processing of ascending proprioceptive inputs. A subset of CFA ET neurons was jointly modulated by passive muscle stimulation and reach-to-consume perturbation (**Fig. 4E**), demonstrating the integration of muscle-specific proprioceptive feedback. Passive forelimb muscle stimulations evoked significant responses in 37.8% of CFA ET neurons (211/558; **Fig. S5F**), 71.1% of which (150/211) were also activated during reach-to-consume. Both passive-responsive (211/558) and passive-non-responsive (347/558) groups contained neurons with brief or sustained reach-to-consume activity patterns, with passive-responsive neurons showing slightly earlier onset of calcium activity (**Fig. S5F**). Perturbation effects were observed in 55.9% of passive-responsive and 57.1% of passive-non-responsive neurons, with larger mean effects in passive-responsive neurons (**Fig. 4H**). A linear model incorporating reach activity, passive responses, and their interaction revealed that in 53.4% of neurons (298/558), perturbation effects were not explained by linear summation, as interaction terms accounted for significant variance (**Fig. S5G**). Together, these results indicate that ET population responses likely incorporate proprioceptive feedback under both natural and perturbed conditions.

Do CFA ET dynamics also integrate the prediction of perturbed proprioceptive feedback? If so, their perturbation-specific activity should decline as the perturbation becomes predictable across trials. To test this, we analyzed the perturbation effects in a session-wise multidimensional activity space, where each axis corresponds to the activity of a single neuron (**Fig. 4I**). We identified a **Perturbation Discriminant Axis (PDA)** that maximally separates perturbed from unperturbed trials for each session. Specifically, we trained a linear discriminant analysis (LDA) model on a random 50% subset of trials to find the PDA, projected the held-out 50% of trials onto the normalized LDA vector and averaged across 10 folds to obtain an unbiased estimate of the perturbation axis for each session. PDA projections of CFA ET dynamics during perturbed reaches diverge sharply from unperturbed ones during movement (**Fig. 4I**). For each session, we quantified a population perturbation effect as the difference between perturbed and control PDA projections. Averaged across sessions, the perturbation effect rose during movement and peaked about 0.5 s after reach onset (**Fig. 4J**). Its amplitude exhibited a modest decay across successive perturbation trials, suggesting neural adaptation along the perturbation discriminant axis (**Fig. 4K**). In line with this population trend, 66/558 (11.8%) neurons changed their activity significantly across perturbation trials (**Fig. 4L**). Control analyses indicated that this decay was not due to shifts in population coding into orthogonal population dimensions (**Fig. S5H**). Together, these results suggest a slight, trial-to-trial attenuation of CFA ET perturbation signals as the perturbation becomes predictable, paralleling the modest behavioral adaptation.

### Neural Dynamics in CFA Multiplex Proprioceptive Perturbations with Motor Output

Transient proprioceptive manipulations rarely abort the progression of reach-to-consume tasks, yet information about motor plans and proprioceptive feedback coexists in ET CFA neurons. We therefore hypothesized that perturbed CFA dynamics are organized into separable subspace. To test this, we quantified the evolution of ET population dynamics in a multidimensional neural activity space.

To visualize the dominant structure of ET activity, we applied principal component analysis (PCA) to the deconvolved, trial-averaged activity of all recorded neurons. The first three principal components (explaining >85% of total variance) revealed that during control reaches, the population trajectory evolved along a C-shaped path after reach onset and remained largely confined to a planar surface (linear regression, R² = 0.92; **Fig. 5A, S6A**). We interpret this plane as a manifold associated with normal reach-to-consume task dynamics. Under proprioceptive perturbation, trajectories diverged from the control shortly after reach onset, and transiently departed from this low-dimensional manifold before gradually returning toward it despite the stimulation remaining active for 2 s (**Fig. 5A**). Moreover, when projected back onto the fitted plane, control and perturbed trajectories traced highly similar geometric paths (Procrustes score = 0.93, **Fig. 5B**), suggesting shared task-related dynamics under the two conditions. Along task-orthogonal directions, however, perturbed trajectories clearly deviated from controls (**Fig. 5C**). This deflection was observed in both passive-responsive and passive-non-responsive ET groups (**Fig. 5D-E**), suggesting that the population-level change might not be solely mediated by ascending sensory signals. These transient departures from the task-relevant manifold indicate that distorted proprioceptive feedback is rapidly integrated into ongoing cortical output dynamics and is primarily encoded along task-orthogonal subspaces.

**Fig 5.**
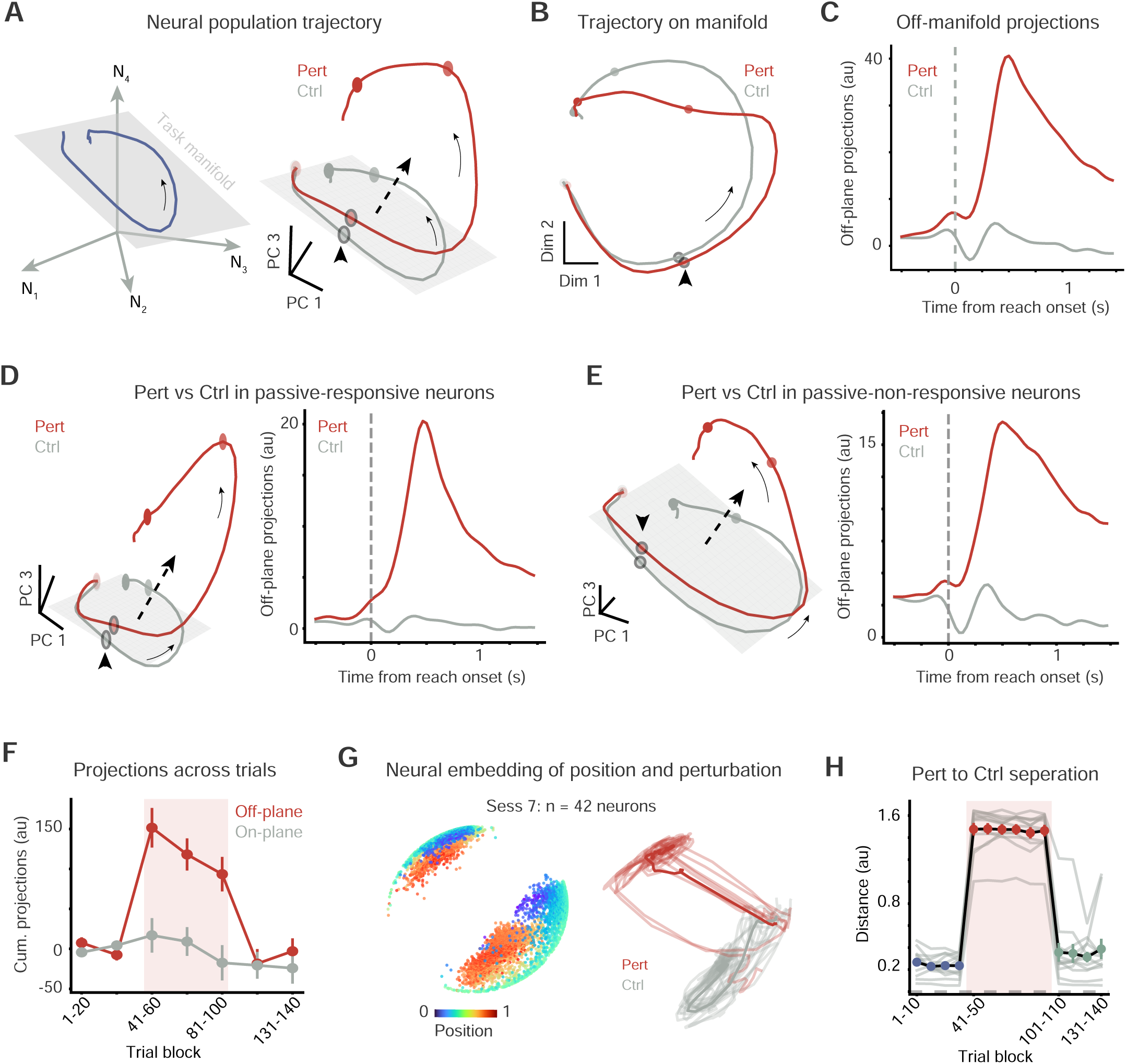
The integration of proprioceptive feedback in CFA ET neuron population dynamics. **(A)** Left: Schematic showing the neural population trajectory on a low-dimensional task-relevant manifold. Right: Normal and perturbed population neural trajectories of CFA dynamics in PC space (n = 558 ET neurons). Arrows indicate the direction of temporal evolution. Circles mark 0.5 s intervals; arrowhead indicate reach onset; plane represents task manifold. **(B)** Normal and perturbed activity projections on the identified task-relevant plane. **(C)** Normal and perturbed task-orthogonal activity projections along the orthogonal direction of the task-relevant plane. **(D)** Comparison of normal and perturbed neural dynamics in passive-responsive ET groups (n = 221 neurons). **(E)** Normal and perturbed neural dynamics in passive-non-responsive ET groups (n = 337 neurons). Note similar multiplexing of proprioceptive input in task-orthogonal dimensions as that of passive-responsive ET groups. **(F)** Cumulative task-relevant and task-orthogonal projections across trial blocks (two-way ANOVA: Group [task-relevant *vs* task-orthogonal], p = 0.004; Trial block, p < 0.0001; Group × Trial interaction, p = 0.0001; n = 12 sessions; mean ± SEM). **(G)** Non-linear neural embedding of hand movement and forelimb stimulation. Left: Color representation of the positions of each time points from an example session. Right: Neural trajectories of individual trials in the same neural embedding. **(H)** Separation between perturbed and normal neural trajectories across trial blocks (one-way ANOVA: Trial block, p < 0.0001; n = 12 sessions; mean ± SEM).

We next asked how distorted proprioceptive feedback, while largely confined to task-orthogonal dimensions, was integrated into ongoing motor dynamics. To do this, we fit linear models to predict hand trajectories (anterior-posterior and dorsal-ventral positions) from trial-averaged neural activity across multiple temporal lags, selecting the optimal lag per session (**Fig. S6B**, median lag = 0.10 ± 0.08 s, cross validated R² = 0.88 ± 0.06, n = 12 sessions). In each session, the first two dimensions of the regression coefficient matrix captured over 99.9% of its variance, suggesting that only two dimensions in neural activity space were capable of predicting behavior, while the rest were output-null. Critically, despite the fact that proprioceptive perturbations were encoded along dimensions orthogonal to the task manifold, they were *not* orthogonal to the prediction manifold (**Fig. S6C**), indicating that proprioceptive disturbances are not strictly confined to null subspaces but partially interact with movement-encoding components.

Finally, we examined whether the off-manifold deflection adapts across repeated stimulation exposures, as suggested by our earlier analysis. We analyzed off-plane projections of neural activity averaged across trial blocks and found a slight trend of attenuation across consecutive perturbation blocks in 8/12 sessions (**Fig. 5F**), paralleling gradual behavioral stabilization. On average, the off-plane projections across perturbation blocks only show a modest attenuation trend (linear regression, *p* = 0.055). Consistent results were obtained using supervised nonlinear contrastive embedding for behavior and representation analysis (CEBRA). Multisession neural embeddings were trained jointly on kinematics and stimulation labels across sessions (**Fig. S6D**). The same CFA ET population reliably codes both task-relevant and perturbation-related signals (**Fig. 5G**). In single-trial embeddings, transient perturbations consistently deflected trajectories away from controls (**Fig. 5G, S6E**). No significant trend in CEBRA separation was observed across perturbation trials (**Fig. 5H**, linear regression, p = 0.42). The minor or absence of a decreasing trend is consistent with perturbation-related dynamics occurring primarily in dimensions orthogonal to task-relevant manifolds, allowing stable behavior despite repeated distortions.

Together, these findings reveal that cortical populations use task-orthogonal dimensions to encode distorted proprioceptive feedback while also allowing this feedback to drive changes in ongoing motor dynamics, potentially preserving both sources of information for readout by other downstream areas.

## Discussion

Precise forelimb control relies on continuous monitoring of proprioceptive feedback. Selective manipulation of muscle-specific proprioceptive input during movement is necessary to uncover how such feedback shapes cortical dynamics. By transiently perturbing forelimb signals during a mouse reach-to-consume task while recording neuron type-specific activity, we show that distorted proprioceptive feedback is integrated into ongoing motor output dynamics with minimal impact on task-relevant activity. In the caudal forelimb area (CFA), individual ET neurons encode both movement kinematics and proprioceptive feedback, and their activity contributes to stabilizing evoked kinematic deviations. At the population level, ET activity rapidly integrates proprioceptive feedback in subspaces largely orthogonal to the task manifold, limiting the influence of perturbation-related signals on evolving task dynamics and safeguarding behavior.

Consistent with the notion that cortical populations implement distinct computations by partitioning activity into separable subspaces ^32,33,42^, our results indicate that distorted proprioceptive feedback is primarily integrated into task-orthogonal dimensions, thereby preserving feedback information while maintaining stable task dynamics within the same ET population. Although our recordings focused specifically on CFA ET output neurons, the C-shaped trajectories observed during normal reaches parallel the rotational population dynamics described in primate and human motor cortex ^48,49^. Cortical recordings have revealed diverse and distributed activity patterns during mouse reach-to-consume behavior ^37,44,50–52^, but how proprioceptive feedback is embedded in cortical output channels has been less clear. In our forelimb muscle stimulation paradigm during the reach-to-consume task, transient activity deflections under proprioceptive distortion indicate rapid integration of feedback within the ET output channel. These transient subspace deflections resemble, but are mechanistically distinct from, the long-term reconfigurations of null dimensions reported during visuomotor and force adaptation in primates ^40,41,53–56^, suggesting that distorted proprioceptive feedback is handled through rapid modulation of task-orthogonal subspaces rather than remapping of the task manifold.

We leveraged optogenetic muscle stimulation to selectively manipulate proprioceptive signals with minimal off-target activation, pain, and muscle fatigue ^57,58^. Although optogenetic muscle stimulation has been previously explored ^59–62^, we achieved robust, temporally precise, and muscle-specific stimulation of the moving forelimb by combining a muscle-enriched opsin expression with a subcutaneous LED implant. Under our protocol (10 Hz, 40 mW, 2 s), local temperature increased only slightly (< 2 °C), and we observed no nocifensive behaviors. This approach therefore allows us to preferentially distort proprioceptive somatosensory signals during natural reach-to-consume behavior, in contrast to magnetic force fields that constrain movements to planar push-pull actions. Compared with sensory nerve stimulation ^63,64^, the forelimb displacement-return cycle evoked by optogenetic muscle stimulation in passive preparations is largely explained by intrinsic muscle contraction-relaxation mechanics and does not appear to rely on short- or long-latency reflexes ^65,66^.

Across perturbed trials, both behavioral and neural adaptation were modest, and we observed no aftereffects when forelimb stimulation was removed. This differs from classical force-field paradigms, where extended exposure produces robust learning and pronounced aftereffects that recruit cerebellar ^67,68^ and cortical circuits ^29,69^. This difference likely reflects that our optogenetic perturbation is not tightly coupled to movement kinematics, unlike traditional force fields, a coupling that appears critical for consolidating long-term internal-model-based learning ^8,27^. Trial-dependent attenuation in a subset of ET neurons, together with declining population activity along a perturbation-discriminant axis, echoes classic proposals that increased motor cortex discharge under external load reflects mismatches between intended and executed movement ^17,70,71^.

Proprioceptive processing engages widely distributed brain structures. From spinal and brainstem circuits, second-order sensory neurons broadly distribute proprioceptive signals to the cerebellum via dorsal and cuneocerebellar pathways and to thalamus and sensorimotor cortex via the dorsal column-medial lemniscal pathway ^2–4^. Rodent studies have often emphasized somatosensory cortex, which shows selective modulation by tactile and proprioceptive inputs and behavioral deficits after silencing ^29,72–75^, whereas primate and human work implicates broader motor areas and cerebellar networks ^8,16,19–21,68,69,76^, highlighting distributed processing of proprioceptive forelimb signals. In our mouse reach-to-consume task with muscle stimulation, the evoked deviation-and-return pattern likely reflects elastic limb mechanics shaped by active CFA modulation. Within this distributed architecture, CFA processes muscle-derived proprioceptive signals to stabilize disturbed forelimb movements (**Fig. 3**), likely by selectively modulating agonist and antagonist muscle tone via descending output ^77^. We therefore propose that ongoing CFA output modulates subcortical and spinal circuits that control forelimb biomechanics to stabilize stimulation-evoked deviations, a hypothesis that will require further investigation.

In summary, using precisely timed optogenetic stimulation of forelimb muscles during a mouse reach-to-consume task, we show that CFA ET neurons integrate distorted proprioceptive signals from muscles with descending dynamics to stabilize perturbed forelimb movements, and segregate perturbation-related activity into dedicated subspaces that preserve task-relevant dynamics for downstream readout.

## FIGURE LEGEND

**Fig S1.**
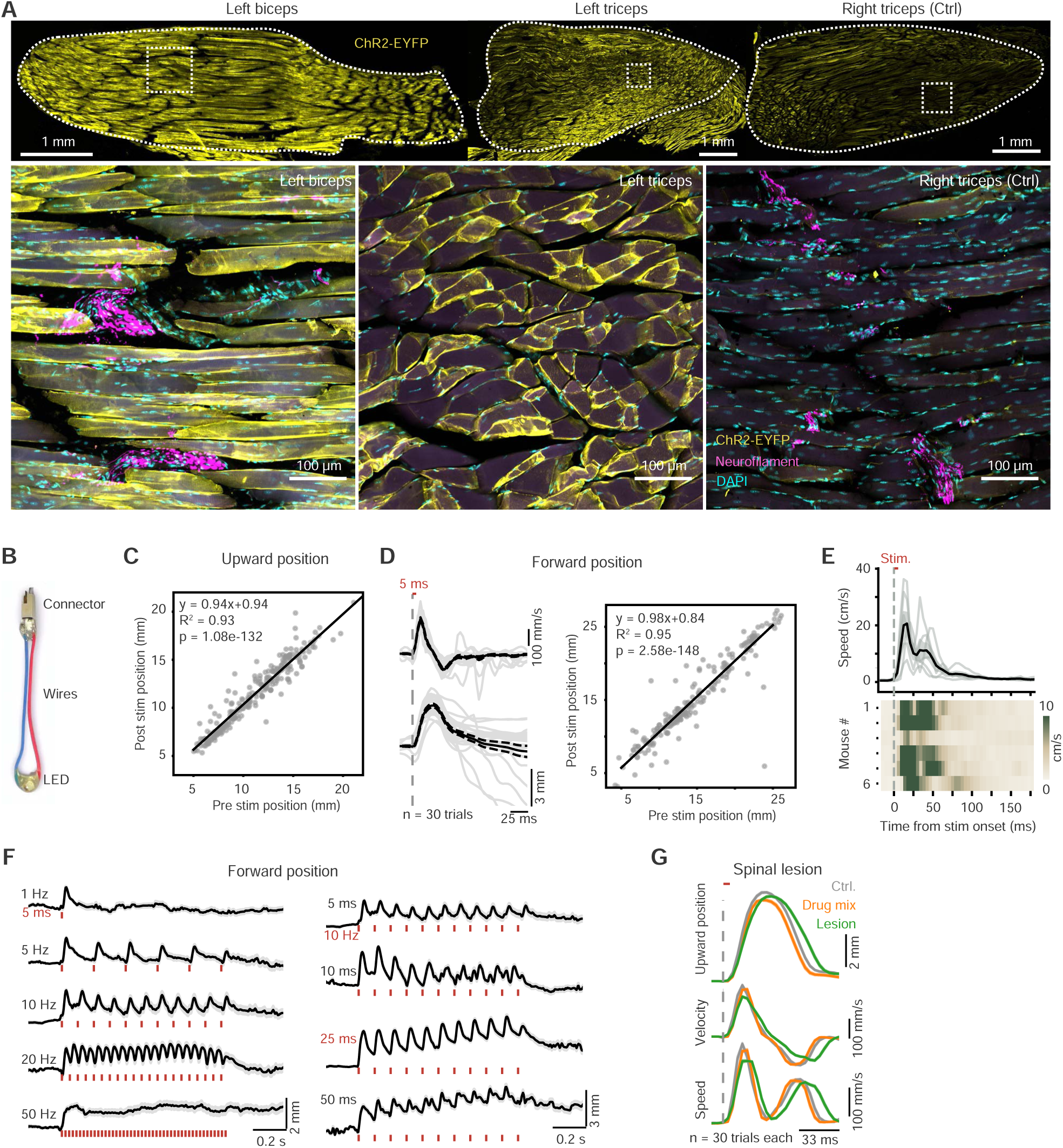
Optogenetic stimulation of forelimb muscles. Related to Fig 1. **(A)** Confocal sections of triceps and biceps muscles stained with GFP (green) and neurofilament light (NFL, red) antibodies. ChR2-EYFP virus was injected into the left forelimb; the right triceps served as control. NFL signals mark nerve endings around intrafusal fibers. Bottom panels show zoomed-in views of dashed regions in the top panels. **(B)** The optogenetic implant. **(C)** Linear regression of hand upward-downward position before versus after stimulation shows that the limb reliably returned to its original position following brief upward displacement (n = 232 trials). **(D)** Left: Example forward-backward hand kinematics evoked by a single 5 ms light pulse from a session (n = 30 trials). Right: Linear regression of forward-backward hand position before versus after stimulation confirms reliable return after brief displacement (n = 232 trials). **(E)** Summary of single 5 ms light pulse-evoked speed changes (n = 6 mice, mean ± SEM). **(F)** Forward forelimb movement evoked by 1 s trains of light pulses across frequencies and pulse durations (n = 6 mice, mean ± SEM). **(G)** Spinal silencing with drug mix or lesion on the evoked movement kinematics in an anesthetized animal (n = 30 trials each).

**Fig S2.**
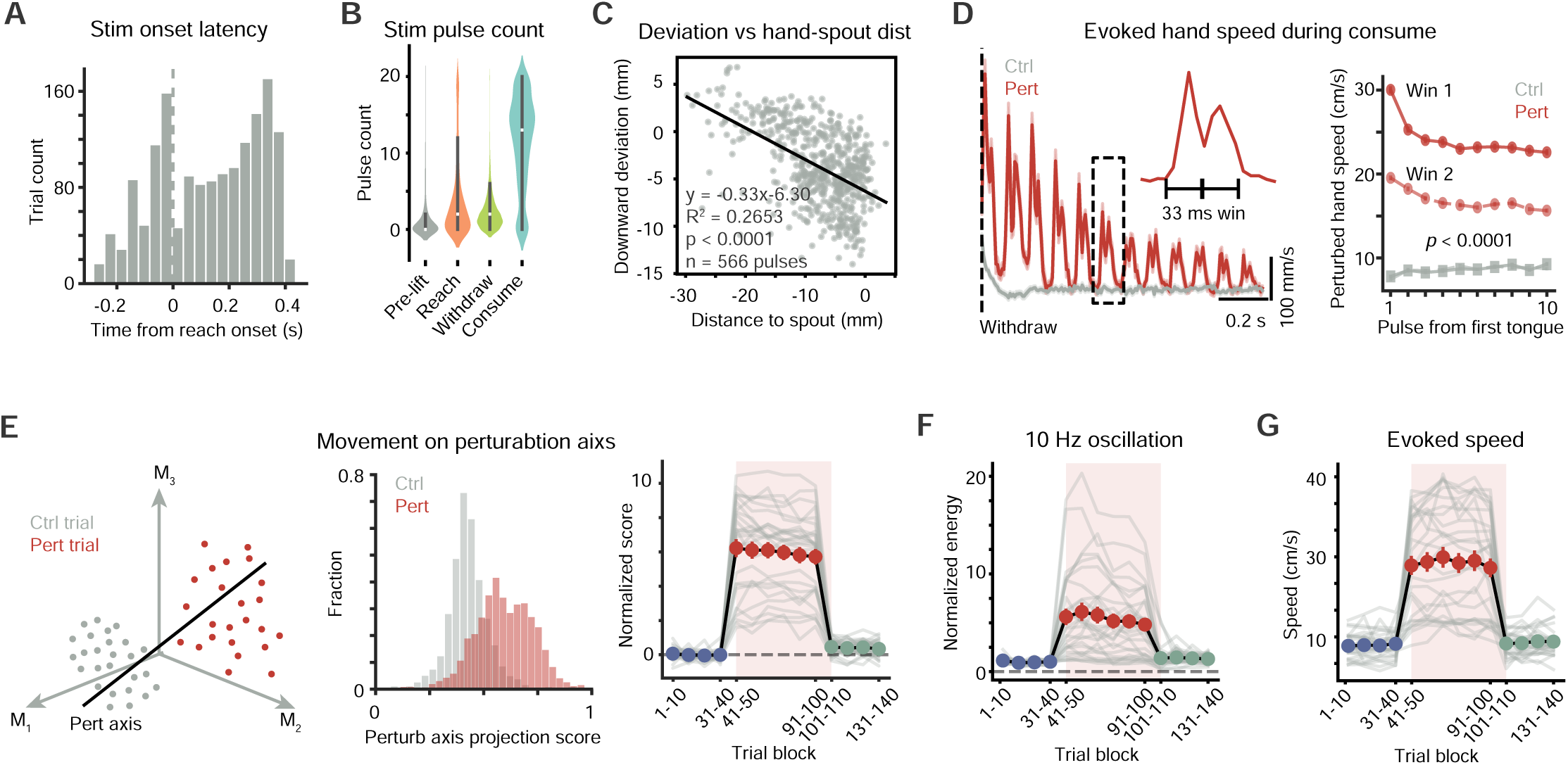
Deviated forelimb movements during reach-to-consume. Related to Fig 2. **(A)** Distribution of stimulation onset latency relative to the reach onset (n = 1582 trials). **(B)** Forelimb stimulation pulse counts during different action phases of reach-to-consume behavior (n = 872 trials). **(C)** Downward deviation amplitude upon perturbation increases as the hand approaches the waterspout during reaching (n = 566 pulses). **(D)** Left: Average hand speed aligned to reach endpoints (n = 80 Ctrl and 60 Pert trials; mean ± SEM). Inset: two speed peaks evoked by each perturbation pulse. Right: Perturbation-evoked peak movement speed during two consecutive windows after each pulse onset was significantly higher than that during virtual stimulations (two-way ANOVA for each window: Group [Ctrl vs Pert], p < 0.0001; Pulse sequence, p < 0.0001; Group × Pulse interaction, p < 0.0001; n = 1620 Ctrl and 1620 Pert trials; mean ± SEM). The first stimulation pulse after tongue protrusion in each trial was used as reference for pulse alignment. **(E)** Left: Identification of a behavior discriminant axis that maximally separated perturbed from unperturbed movements in multidimensional kinematic space. Middle: Distribution of kinematic projection scores on the discriminant axis (Wilcoxon rank-sum test, p < 0.0001; n = 1620 Ctrl and 1620 Pert trials). Right: Projection scores across trials of each session (n = 27 sessions; mean ± SEM). Scores were averaged per 10 trials and baseline-subtracted for each session. **(F)** Oscillatory movement kinematics across trials (n = 27 sessions; mean ± SEM). **(G)** Evoked hand speed across trials (n = 27 sessions; mean ± SEM).

**Fig S3.**
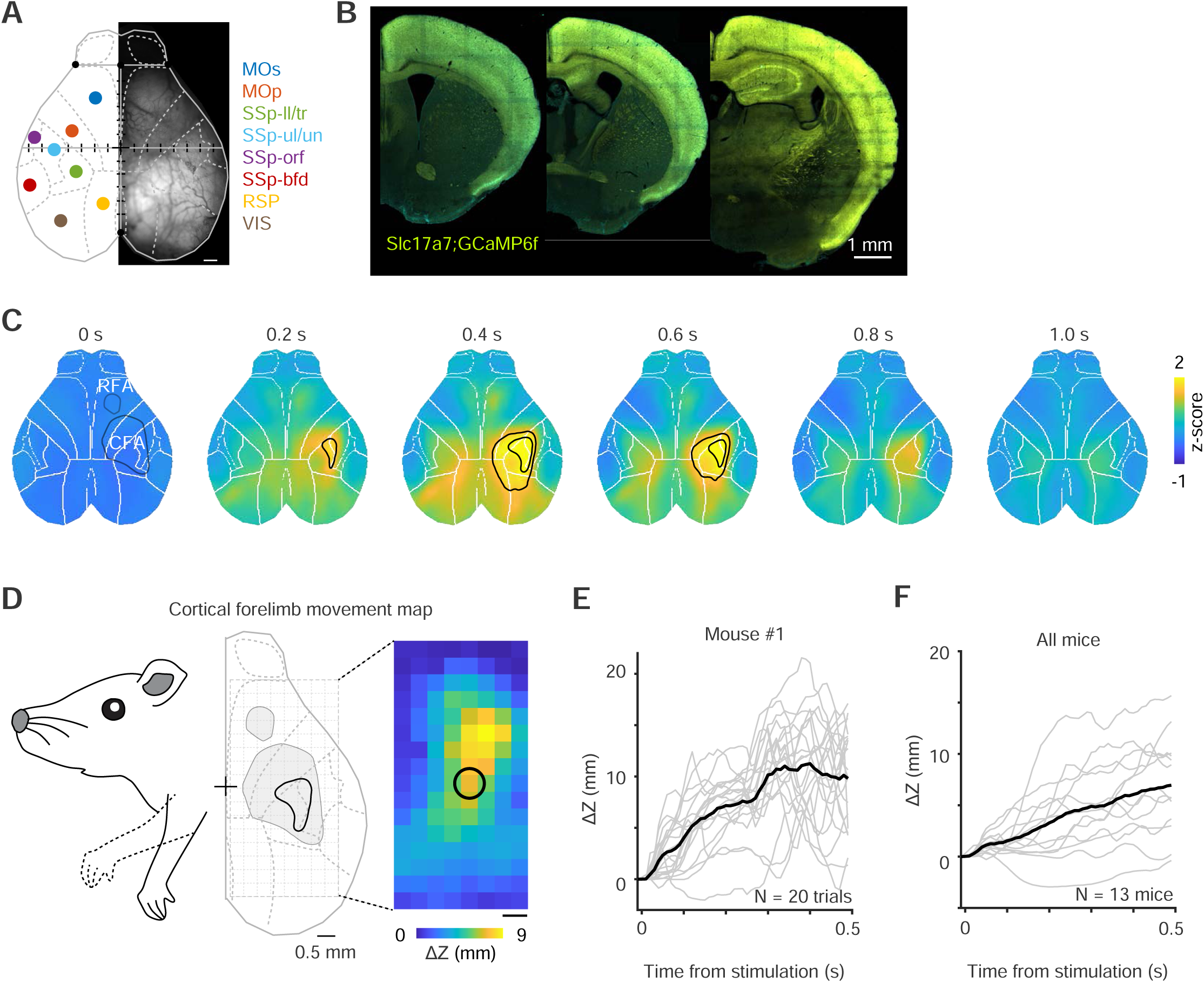
Forelimb muscle stimulation preferentially activates caudal forelimb area. Related to Fig 3. **(A)** Registered atlas regions overlaid on a dorsal view of mouse cortex. MOB, main olfactory bulb; MOs, secondary motor cortex; MOp, primary motor cortex; RSP, retrosplenial cortex; SSp, primary somatosensory cortex; tr, trunk; ll, lower limb; ul/un, upper limb/unknown region; orf, orofacial; bfd, barrel field; VIS, visual cortex. **(B)** Coronal sections showing cortical projection neurons in *Slc17a7-ires-Cre;Ai148* mice. **(C)** Average cortical activity image under passive forelimb muscle activation across time (n = 6 sessions). Caudal and rostral forelimb areas are indicated. **(D)** Heatmap of forelimb movement evoked by optogenetic activation of extratelencephalic neurons in different cortical areas (n = 13 *Fezf2-CreER;Ai32* mice). The heatmap shows upward displacement during 0.5 s of stimulation (5 ms pulses at 50 Hz). Each square corresponds to a stimulation site (375 µm). **(E)** Example forelimb displacement traces during stimulation at the region of interest (circled site in D, n = 20 trials). **(F)** Summary of displacement across all mice (n = 13, mean ± SEM).

**Fig S4.**
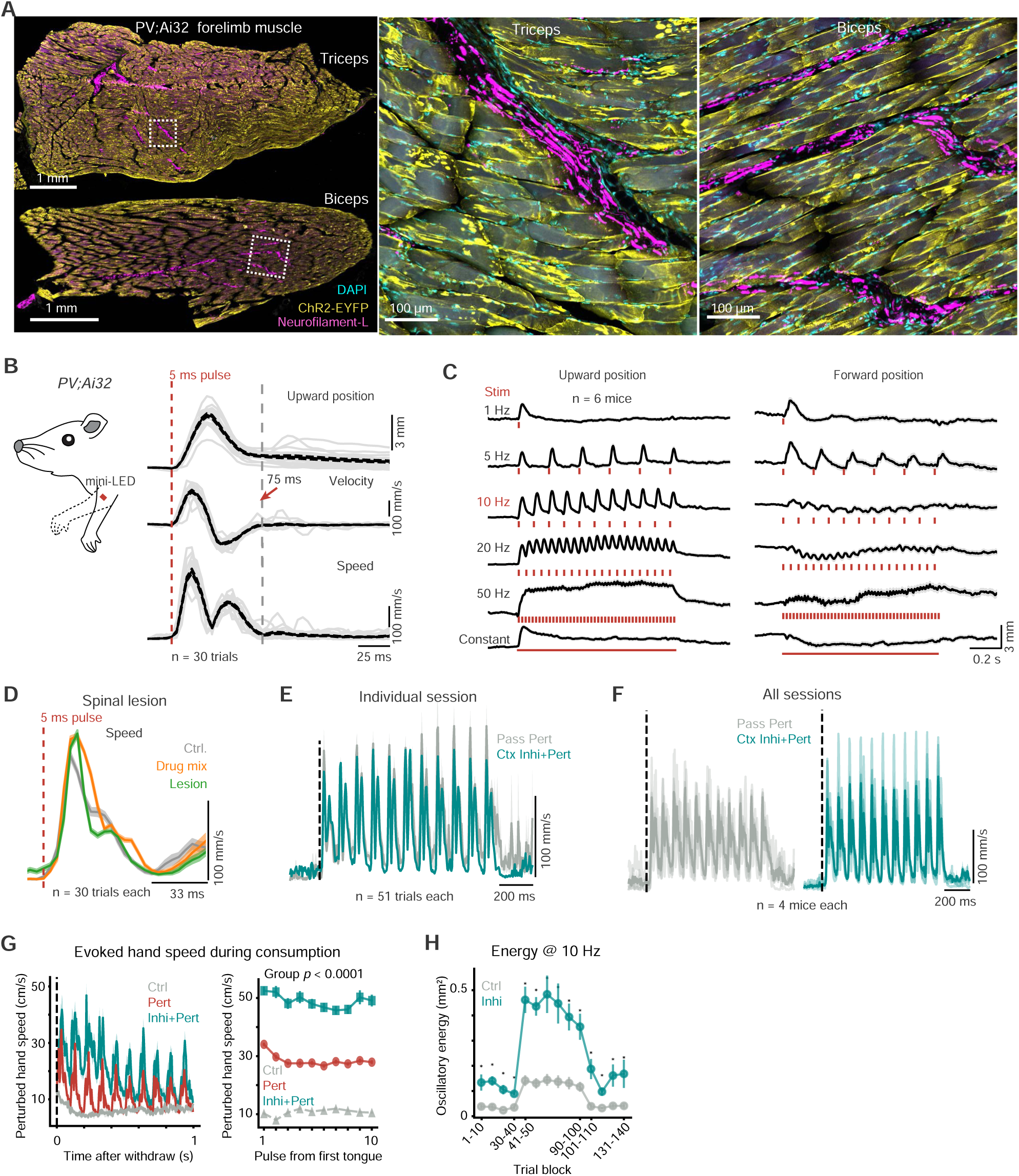
Inactivating cortex during forelimb perturbation. Related to Fig 3. **(A)** ChR2-EYFP expression in *PV;Ai32* mouse forelimb muscles. Confocal sections of triceps and biceps muscles stained with GFP and neurofilament light antibodies. **(B)** Hand movement kinematics evoked by a single 5 ms light pulse from a session (n = 30 trials). **(C)** Upward and forward forelimb movement evoked by 1 s trains of 5 ms light pulses at 1 Hz, 5 Hz, 10 Hz, 20 Hz, 50 Hz and constant (n = 6 mice, mean ± SEM). **(D)** Spinal silencing or lesion on the evoked movement kinematics in an anesthetized *PV;Ai32* animal (n = 30 trials each). **(E)** Inactivating CFA on evoked movement speed from an example session of *PV;Ai32* animal (n = 30 trials each). **(F)** Summary of the effect of inactivating CFA on evoked movement speed in resting *PV;Ai32* mice (n = 4 mice). **(G)** Left: Average hand speed aligned to reach endpoints (n = 80 Ctrl, 60 Pert and 60 Inhi+Pert trials; mean ± SEM). Inset: two speed peaks evoked by each perturbation pulse. Right: Perturbation-evoked peak movement speed for each pulse (two-way ANOVA: Group [Pert vs Inhi+Pert], p < 0.0001; Pulse sequence, p < 0.0001; Group × Pulse interaction, p < 0.0001; n = 1120 Ctrl, 840 Pert and 240 Inhi+Pert trials; mean ± SEM). **(H)** Cortical silencing on oscillatory movement kinematics across trials (two-way ANOVA: Group [Ctrl vs Inhi], p < 0.0001; Trial block, p < 0.0001; Group × Trial interaction, p < 0.0001; Bonferroni post hoc test; n = 14 Ctrl and 4 Inhi sessions; mean ± SEM).

**Fig S5.**
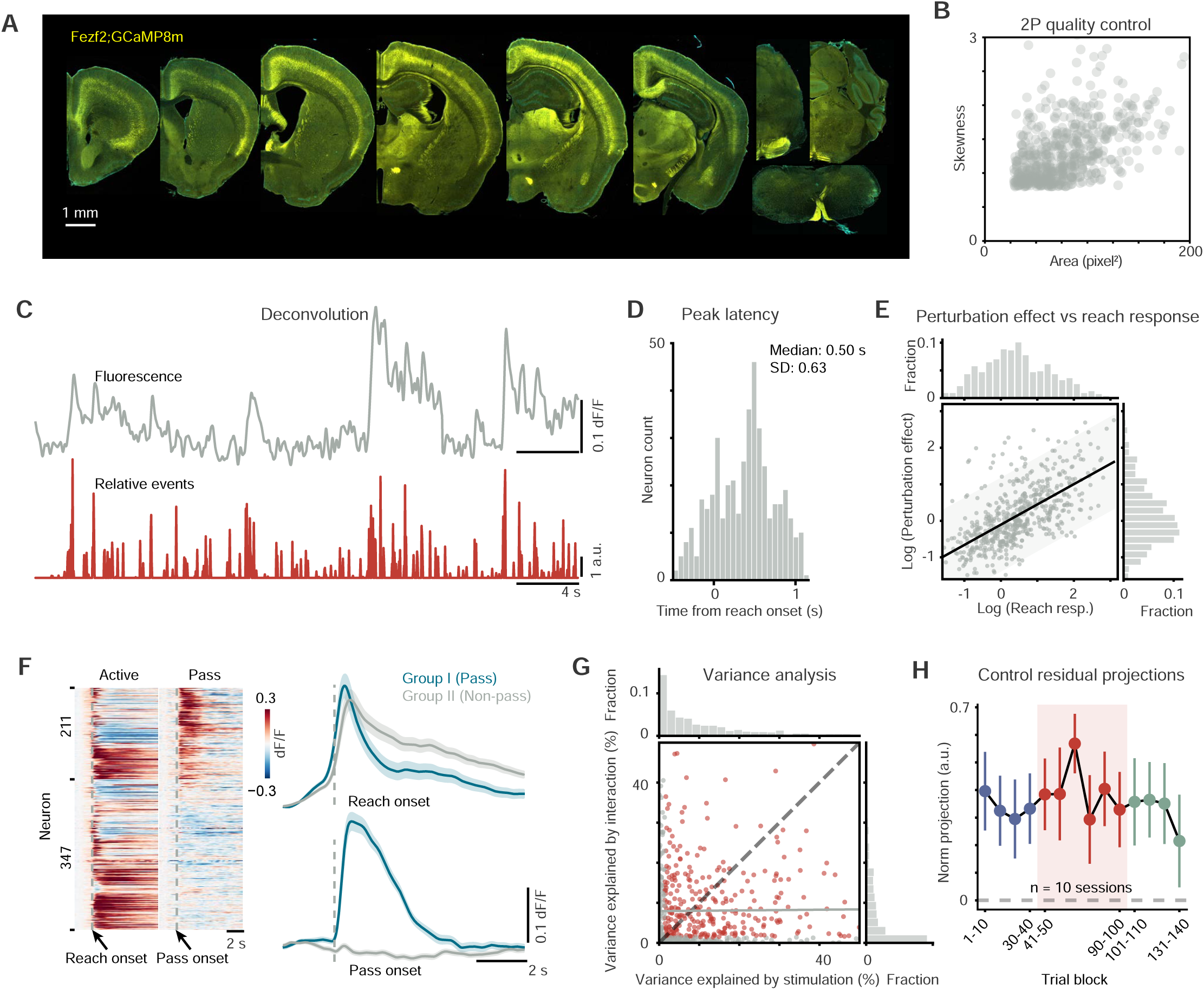
Two-photon imaging of CFA ET neurons. Related to Fig 4. **(A)** Coronal brain sections showing GCaMP expression in cortical extratelencephalic neurons in *Fezf2-CreER;GCaMP8m* mice. **(B)** Quality control of two-photon data based on skewness of activity and areas of individual neurons (n = 558 neurons). **(C)** Deconvolution of calcium fluorescence signals into transient spike events. **(D)** Peak latency distribution of the perturbation effect (n = 558 neurons; median ± SD = 0.50 ± 0.63 s). **(E)** Dependence of perturbation effect on reach-to-consume activity amplitude (R² = 0.395, n = 558 neurons). Each dot represents an individual neuron. Histograms show the distributions of reach response amplitude (top) and perturbation effect (right). **(F)** Left: Heatmap summary of individual neuronal responses to reach-to-consume and passive muscle activation. Right: Comparison between Group I (passive-responsive, n = 211 neurons) and Group II (passive-non-responsive, n = 347 neurons). Group I neurons exhibited earlier rise and faster decay kinetics than Group II (mean ± SEM). **(G)** Variance explained by reach-to-consume activity, passive muscle evoked activity, and their interaction in a mixed linear model. Significant interaction effect in 391/558 neurons. **(H)** No decay along LDA orthogonal dimensions across trials of perturbation block (n = 10 sessions).

**Fig S6.**
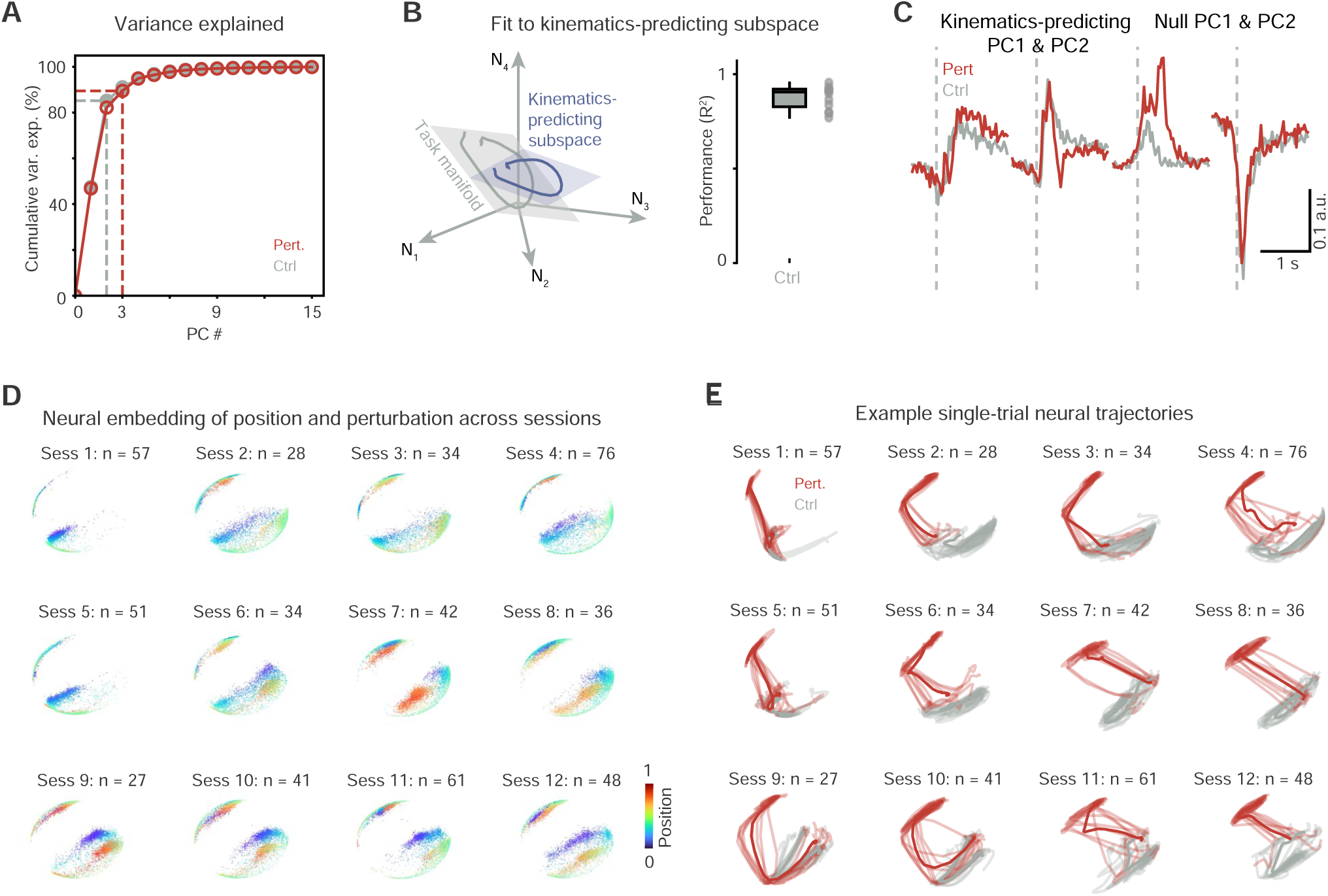
Neural population dynamics. Related to Fig 5. **(A)** Cumulative variance explained by the principal components of averaged normal and perturbed activity (n = 558 ET neurons). **(B)** Left: Schematic showing identification of the kinematics predicting subspace and task-manifold. Right: Summarized performance of linear repression in identifying kinematics-predicting subspaces (n = 12 sessions). Box plot: center line, median; box limits, 25th-75th percentiles (interquartile range, IQR); whiskers, 1.5× IQR. **(C)** Comparison of the projections of perturbed and normal neural population activity onto the kinematics-predicting and null dimensions from an example session. **(D)** Non-linear CEBRA neural embedding of hand movement and perturbation. Heatmap indicates the positions of each time point. **(E)** Example single trial neural trajectories of perturbed and normal neural trajectories in a multi-session behavior supervised CEBRA embedding (n = 10 trials each).

## RESOURCE AVAILABILITY

### Animals and materials availability

All mice, reagents and materials are openly available with detailed source information listed in the Methods.

### Data and code availability

Data and code will be made publicly available upon publication.

## ACKNOWLEDGEMENTS

We thank Dr. Grimm for sharing the AAVMYO helper plasmids; Drs. Herzfeld and Whishaw for helpful comments and discussions; Dr. Zhong for schematic illustrations; and Baoxia Han for animal care. This work was supported by the NIMH research project U19MH114823-01 grant to Z.J.H. Z.J.H. is also supported by an NIH Director’s Pioneer Award 1DP1MH129954-01.

## AUTHOR CONTRIBUTIONS

Y.L. conceived the study, designed experiments, built setups, collected data, and established data-analysis pipelines. X.H.X performed surgeries and, with Z.Q., trained mice for two-photon imaging. J.P. and S.P. conducted computational modeling. X.A. performed motor mapping and spinal lesion surgeries. S.Z. packaged viral vectors. Y.L. and P.J.M explored forelimb muscle injections. Z.J.H provided funding and supervised the project. Y.L. Z.J.H., J.P. and N.L. wrote the manuscript with edits from all authors. All authors discussed and interpreted the data.

## DECLARATION OF INTERESTS

The authors declare no competing interests.

### Methods Mice

Animal care, use, surgical and behavioral procedures conformed to the guidelines of the National Institutes of Health. The experiments were approved by the Institutional Animal Care and Use Committee of Duke University. Experiments were conducted with 8-week- to 16-week-old male and female mice. The number of animals used in each experiment is noted in the corresponding section. Mouse strains were: *Slc17a7-IRES2-Cre-D* (B6.Cg-Slc17a7^tm1.1(cre)Hze^/J, RRID:IMSR_JAX:037512), *Ai148D* (B6.Cg-Igs7^tm148.1(tetO–GCaMP6f^,^CAG–tTA2)Hze/^J; RRID:IMSR_JAX:030328), *Fezf2-CreER* (B6;129S4-Fezf2^tm1.1(cre/ERT2)Zjh^/J; RRID:IMSR_JAX:036296), *TIGRE2-jGCaMP8m-IRES-tTA2-WPRE* (*Igs7^tm1^*^(tetO–GCaMP8m,CAG–tTA2)^*^Genie/^J*; RRID:IMSR_JAX:037718), *Pvalb-IRES-Cre* (B6.129P2-Pvalb^tm1(cre)Arbr^/J; RRID:IMSR_JAX:017320), *Ai32* (B6.Cg-Gt(ROSA)26Sor^tm32(CAG–COP4*H134R/EYFP)Hze^/J; RRID:IMSR_JAX:024109). Mice were housed in groups of up to five mice per cage, in a room with a 12/12 light/dark cycle. After surgery, mice were housed in a new home cage individually or with familiar groups for at least one week prior to further experiments.

### Surgery

Surgery was performed as previously described. For head-restrained experiments, a titanium flat headpost was secured on the skull with dental cement (C&B Metabond, Parkell; Ortho-Jet, Lang Dental). For wide-field calcium imaging, the skull with most of the dorsal cortex was exposed, cleaned with saline and coated with a thin layer of cyanoacrylate glue (Zap-A-Gap CA+, Pacer Technology) to render the bone transparent. For cortical silencing in *Pvalb-IRES-Cre;Ai32* mice, a thin-skull preparation was covered with low-toxicity silicone adhesive (KWIK-SIL, World Precision Instruments) to protect the cortex. For two-photon imaging, a craniotomy was made and a two-layer 3 mm cranial window was implanted over the sensorimotor cortex.

To perturb forelimb movement, viral vector injections into forelimb muscles were performed using insulin needles. A total of 6 μL of *AAVMYO3-CAG-DIO-hChR2-EYFP* (3.2 × 10¹³ vg/mL) was diluted in 54 μL of sterile saline, and six injections (10 μL each) were delivered into the proximal muscles of the left forelimb (including triceps and biceps). After 21 to 28 days of expression, an optogenetic implant was placed subcutaneously in the left forelimb. The implant consisted of a mini-LED (LXZ1-PB01, LumiLEDs), two thin flexible wires, and an electronic connector (A79604-001, Omnetics). The LED was encapsulated in a thick layer of clear epoxy and tested in saline to verify insulation. The final implant weight was ∼0.150 g. For implantation, a small incision was made in the dorsal neck, and the LED with attached wires was tunneled subcutaneously toward the proximal forelimb muscles. The incision was closed along the wire using tissue adhesive, and the connector was secured to the skull with dental cement.

### Tamoxifen induction

Temporal control of Cre recombinase was achieved via intraperitoneal tamoxifen injections (2 × 100 mg/kg, 20 mg/ml in corn oil) in *Fezf2-CreER*; *jGCaMP8m* mice. The first induction occurred at weaning and the second one week later.

### Immunohistochemistry

After the experiments, animals were euthanized with isoflurane and transcardially perfused with saline, followed by fixation with 4% paraformaldehyde. Forelimb muscles (triceps and biceps) were dissected, fixed, sectioned at 20 µm thickness with cryostats, and mounted onto slides. Immunohistochemistry was performed on the mounted muscle sections. Briefly, sections were pretreated in blocking solution 10% Blocking One in PBS containing 0.3% Triton X-100) for 1 h at room temperature, followed by overnight incubation at 4 °C with primary antibodies diluted in the same blocking solution. The next day, sections were washed in blocking solution and incubated with secondary antibodies for 2 h at room temperature. Confocal images were acquired using a ZEISS Axio Observer microscope equipped with an LSM 900 laser system.

### Passive muscle stimulation

Mice with optogenetic forelimb implants were head-restrained, with the lower body supported in a tube and both forelimbs hanging freely. Animal forelimb movement was videotaped under infrared illumination with two synchronized high-speed (240 fps) cameras from front and side views. Videos were acquired using a custom Bonsai workflow and analyzed offline. The duration and frequency of optogenetic muscle stimulation via mini-LED were controlled by customized MATLAB scripts and synchronized with video acquisition.

To examine the spinal contribution to passively evoked forelimb movements, experiments were conducted in anesthetized mice under isoflurane. Three consecutive sessions (baseline, drug application, and mechanical lesion) of optogenetic stimulation were performed. Each session consisted of 30 stimulation trials over ∼5 min. A cocktail of synaptic blockers (200 µL) targeting glutamatergic (5 mM DNQX, 1 mM AP5) and GABAergic (1 mM gabazine) transmission was applied to the exposed cervical spinal cord, followed by a 10 min wait period before stimulation. After drug application, a mechanical lesion of the exposed cervical spinal cord was made using a surgical blade. Finally, mice were euthanized, and the forelimb was excised and immersed in saline for *ex vivo* forelimb stimulation.

### Mouse reach-to-consume task

The day before behavioral training, mice were weighed and transferred to a new cage with restricted water access, receiving supplemental water each day to maintain >80% of their initial body weight. The reach-for-water task was controlled in real time using custom MATLAB code, interfaced with piezo sensors, water valves, and linear actuators via a data acquisition board (USB-6351, National Instruments). Mice were trained to grasp sucrose solution drops (10% w/v, 20-50 μL) delivered in the peri-orofacial space, with the waterspout centered and fixed in front of the nose (∼4 mm below the nose tip). Training consisted of one session per day for three days, with 100 trials per session and random inter-trial intervals of 12-20 s. Following training, a typical experiment session comprised 40 pre-perturbation baseline, 60 perturbation, and 40 post-perturbation recovery trials arranged in blocks. During the perturbation block, 10 Hz light pulses (5 or 25 ms) were delivered around reach onset to induce optogenetic forelimb perturbations. No stimulation was applied during baseline or recovery trials. Animal behavior was videotaped under infrared illumination at 240 fps.

### Inactivating cortex

Cortical inactivation was performed in *Pvalb-IRES-Cre;Ai32* mice via optogenetic activation of interneurons. Blue light pulses (473 nm, 5 ms, 50 Hz) were delivered on 50% of reach trials during perturbations. A 200 μm fiber tip contacted the thinned skull, positioned with micromanipulator, and light intensity was set to 10 mW at the fiber tip. Photoinhibition light was delivered from 100 ms before perturbation onset to 100 ms after perturbation offset in perturbation trials.

### Cortex-wide calcium imaging

Cortex-wide calcium imaging was performed using an inverted tandem-lens macroscope (105 mm top, 85 mm bottom; magnification ×1.24) and an sCMOS camera (Edge 5.5, PCO), yielding a 12.4 × 10.5 mm field of view at ∼20 µm/pixel. GCaMP fluorescence was captured with a 525 nm band-pass filter under alternating blue (470 nm) and violet (405 nm) LED excitation, combined via dichroic and projected with a 495 nm long-pass mirror. Illumination and camera acquisition (50 fps; 25 fps per wavelength) were synchronized with an Arduino. Violet excitation produced non-calcium-dependent fluorescence, enabling regression-based correction of intrinsic signals; analyses used the differential calcium-dependent signal at 25 fps.

### Two-photon imaging

Activity of individual neurons was imaged in behaving mice using a two-photon microscope (Ultima Investigator, Bruker) coupled to a fs-pulse Ti:Sapphire laser (Mai Tai DeepSee, Spectra-Physics). The laser wavelength was tuned to 1040 nm for excitation of the genetically encoded calcium indicator GCaMP. The system was equipped with an 8 kHz resonant galvanometer and a 16 × water-immersion objective (0.8 NA, Nikon) mounted on a 400 µm-range piezoelectric z-drive. GCaMP fluorescence was filtered (525/70 nm bandpass), detected with GaAsP PMTs (Hamamatsu), and amplified before digitization. Images were acquired in single-plane mode at 512 × 512 pixels (2× digital zoom; field of view ∼512 × 512 µm) at ∼30 Hz. Excitation laser power ranged from 150-400 mW at the source, corresponding to 33-76 mW at the objective. Imaging was performed in a trial-by-trial manner, with each trial consisting of 300 consecutive frames. To block possible light contamination from the optogenetic limb LED, imaging lenses were covered with blackout tape. Animal behavior was simultaneously recorded at 150 fps under synchronized two-photon illumination.

### Data processing and analysis

Data processing and analyses were performed with Python (>= 3.9) or MATLAB 2023b (Mathworks), unless otherwise specified.

### Movement time series

Two DeepLabCut 3.0 networks were trained separately on >2000 frames (1155 front, 1196 side) covering all phases of the forelimb reach-to-consume behavior from multiple sessions across different animals, incorporating variations in color, size, head-restraint, illumination, and behavior phase to ensure robustness. Eighteen front-view and 22 side-view keypoints (digits, nose, mouth, tongue, waterspout, water drop) were labeled per frame.

### Position deviations and evoked speed

Forelimb stimulation-evoked kinematic responses were quantified from hand-position time series. For each trial, the stimulation onset frame was used when available, or set to the average for virtual stimulation onset for control trials. The post-onset segment was divided into consecutive pulses of duration equal to the stimulation cycle (100 ms), with each pulse further subdivided into three equal sub-windows. Within each sub-window, positional deviation was computed as the dot product between the displacement vector (final minus initial hand position vector) and a trial-specific reference axis, yielding a 3-D array of range lengths (trial × pulse × sub-window). Instantaneous speeds were calculated as the Euclidean norm of frame-to-frame displacement vectors multiplied by the sampling rate, and the maximum speed per sub-window was extracted as the evoked speed. We use the max positional deviation and the evoked speed in the first sub-window (33 ms) across all pulses in a trial to capture the trial-level deviation magnitude following stimulation.

### Oscillatory energy

Transient 10 Hz oscillations were quantified from hand position time series over 2 seconds following stimulation onset. Each trial’s post-onset segment was divided into five equal non-overlapping windows (400 ms); within each window, linear trends were removed and a Hann taper applied. The one-sided fast Fourier transform was computed, and the power spectral density (PSD) was calculated for each window. The frequency bin closest to 10 Hz was identified, and PSD at that bin was converted to total power (mm²) by multiplying by the bin width. The maximum power at 10 Hz across windows was taken as the trial-level 10 Hz energy metric.

### Linear discriminant analysis on kinematics

To compare forelimb kinematics of perturbation and control trials, we applied linear discriminant analysis (LDA) to features derived from principal component analysis (PCA). PCA was performed on z-scored forelimb kinematic data from both front and side camera views by flattening all trials and time points into a single matrix. For inhibition trials, the analysis window corresponded to the actual forelimb stimulation period. For control trials, a virtual stimulation window of equivalent duration was used to ensure temporal alignment. Within each trial, PCA projections from the analysis window were averaged to produce a single feature vector. LDA was performed on the PC projections using the trial type (perturbation vs. control) as the class label. The LDA axis was determined based on the separation between the two trial types. For each trial, the signed deviation along the LDA axis was computed by projecting the trial’s PC feature vector onto the LDA axis and normalized by subtracting the mean projection of control trials within the same session. We performed 10-fold stratified cross-validation in each session to ensure robustness. The resulting signed LDA deviations were used for visualization, comparisons of animal behavior between trial types, and quantification of the change of perturbed effect across trials.

### Wide-field cortical activity

Cortex-wide calcium dynamics were extracted as described previously ^37^. Landmarks of the dorsal cortex were marked, and a cortical mask was defined from an example frame. Raw videos were split into GCaMP and control channels, cropped, flattened, and concatenated across trials into an n × t matrix (n = 440 × 440 pixels, t = frames). Denoising was performed with singular value decomposition, yielding spatial components (*U*), temporal components (*V*^T^), and singular values (*S*). To reduce computational load, analyses used the compressed representation *V*c = *SV*^T^; results were later reconstructed by multiplying with *U*. This produced a low-rank decomposition *Y*raw = *UV*c + Error, achieving >95% compression with minimal signal loss. Control-channel fluorescence (405 nm) was regressed from GCaMP frames (473 nm) to isolate calcium-dependent signals. Z-score normalization used the 1 s pre-stimulation as baseline. All data were registered to the Allen CCF3 using five anatomical landmarks (left/right olfactory bulb-cortex junctions, retrosplenial base, and Bregma).

### Two-photon neuron activity

Suite2p was used to extract two-photon neural activity ^78^. For soma region of interest (ROI), fluorescence time series traces were produced by averaging the pixels within each ROI for all imaging frames and then subtracting the estimated neuropil fluorescence from them. Neuropil is defined as the signal within a surrounding region around a cell, excluding the cell itself. The neuropil-subtracted fluorescence was used for all further analysis. Only ROIs with size > 28.26 mm^2^ (radius 3 µm) and skewness > 0.8 were good quality neurons and shown in the manuscript.

Change of fluorescence: The baseline (F0) of the fluorescence trace was estimated for each trial from a baseline period (0.5 s before water delivery). The ΔF/F0 was then calculated, where ΔF was first determined by subtracting F0 from the raw trace, and then divided by F0. ΔF/F0 traces were computed for each trial and smoothed with a 0.3 sec moving window.

Relative quantity of spikes: Neuropil-subtracted fluorescence was deconvolved to estimate the time and relative quantity of fluorescent spikes. Non-negative spike deconvolution was implemented using Suite2p with OASIS algorithm under 0.7 s fixed calcium decay. The deconvolved value reflects a relative quantity of spikes rather than the absolute spike rates. We normalized the deconvolved value of each trial by subtracting the average of baseline period.

### Neural population dynamics

The perturbation discriminant axis of neural activity between normal and perturbed trials was defined using principal component analysis (PCA) followed by linear discriminant analysis (LDA). Neural responses from perturbed and unperturbed reach trials (2 s window following stimulation onset) were projected onto principal components, and LDA was applied to extract the coding direction that best separated perturbed from unperturbed trials. The resulting discriminant vector was projected back to neural space and normalized to unit length.

To visualize and quantify the time-evolution of trial-averaged population activity across perturbation conditions, we concatenated deconvolved activity from all neurons into a matrix *X* size *n* × *ct*, where *n* is the number of neurons, *c* the number of perturbation conditions, and *t* the number of time bins (33 ms). Principal component analysis (PCA) was then applied to extract latent dimensions, and neural trajectories were visualized by projecting *X* onto the first three principal components. To identify the low-dimensional task manifold, a plane was fit to the control trajectory (*z = ax + by + c*), and its normal vector was used to compute off-plane task-orthogonal projections, reflecting deviations from the control task manifold during forelimb stimulation. Cross-condition alignment was further quantified by projecting each trajectory onto the control plane and computing Procrustes similarity between conditions.

We estimated the kinematics-predicting and null dimensions by relating trial-averaged neural population activity to simultaneously measured hand kinematics. Neural activity (z-scored spiking) was aligned to hand position and velocity time series, and a temporal lag between neural and behavioral data was optimized per session by multi-output (multivariate) linear regression. The regression weight matrix mapping neural activity to hand kinematics was then decomposed using singular value decomposition (SVD). The column space spanned by the top singular vectors defined the kinematics-predicting subspace (dimensions that linearly predict movement), while the orthogonal complement defined the putative null subspace (dimensions orthogonal to behavior).

We applied nonlinear contrastive embedding for behavior and representation analysis (CEBRA) to uncover low-dimensional neural dynamics linked to behavior ^79^. For each session, CFA calcium activity was aligned to forelimb movement trajectories during reach-to-consume movements. Hand position was computed from digit and paw keypoints, normalized, and resampled to the imaging frame rate (30 Hz). Perturbation periods were labeled, and neural activity and behavioral features were flattened across trials into continuous time series for CEBRA. A multisession model (“offset10-model”) was trained jointly across sessions (>25 neurons; batch size 512; learning rate 3×10⁻⁴; 64 hidden units; 30,000 iterations; cosine distance; temperature = 1). The resulting 3D embeddings captured shared temporal and behavioral structure, enabling visualization of neural trajectories during perturbed and unperturbed reaches. Although the CEBRA space is nonlinear, Euclidean distance provides a local measure of neural state separation relative to the mean normal trajectory. Trial embeddings were analyzed over a 1 s period following reach onset and averaged within 10-trial blocks.

### Statistics & Reproducibility

All experiments were conducted with randomized conditions. Sample sizes were determined with reference to established standards in literature. For each condition, an adequate number of reach-to-grasp trials were acquired to ensure robustness and reproducibility of the results. No statistical method was used to predetermine sample size. No data was excluded from the analysis. Non-parametric statistical tests were used wherever possible. All statistical analyses were performed using two-sided tests.

## SUPPLEMENTARY INFORMATION

**Supplementary Figures S1-S6**

**Supplementary Video 1**

Forelimb movements evoked by optogenetic stimulation of forelimb muscles in a resting mouse, with tracked left-hand keypoints overlaid. Acquired at 240 fps and displayed at 30 fps (1/8 real time).

**Supplementary Video 2**

Mouse reach-to-consume behavior with and without optogenetic muscle stimulation, overlaid with tracked keypoints of the left hand. Videos play at 1/5 real time.

**Supplementary Video 3**

Average widefield calcium movie across 30 trials from a resting *Slc17a7-IRES-Cre;Ai148* mouse during optogenetic stimulation of left forelimb muscles. Stimulation: 0.5 s at 10 Hz. Time 0 ms indicates onset.

